# Tet Controls Axon Guidance in Early Brain Development through Glutamatergic Signaling

**DOI:** 10.1101/2023.05.02.539069

**Authors:** Hiep Tran, Le Le, Badri Nath Singh, Joseph Kramer, Ruth Steward

## Abstract

Mutations in human TET proteins have been found in individuals with neurodevelopmental disorders. Here we report a new function of Tet in regulating *Drosophila* early brain development. We found that mutation in the Tet DNA-binding domain (*Tet^AXXC^*) resulted in axon guidance defects in the mushroom body (MB). Tet is required in early brain development during the outgrowth of MB β axons. Transcriptomic study shows that glutamine synthetase 2 (Gs2), a key enzyme in glutamatergic signaling, is significantly downregulated in the *Tet^AXXC^* mutant brains. CRISPR/Cas9 mutagenesis or RNAi knockdown of Gs2 recapitulates the *Tet^AXXC^* mutant phenotype. Surprisingly, Tet and Gs2 act in the insulin-producing cells (IPCs) to control MB axon guidance, and overexpression of Gs2 in these cells rescues the axon guidance defects of *Tet^AXXC^*. Treating *Tet^AXXC^* with the metabotropic glutamate receptor antagonist MPEP can rescue while treating with glutamate enhances the phenotype confirming Tet function in regulating glutamatergic signaling. *Tet^AXXC^* and the *Drosophila* homolog of *Fragile X Messenger Ribonucleoprotein protein* mutant (*Fmr1^3^*) have similar axon guidance defects and reduction in Gs2 mRNA levels. Interestingly, overexpression of Gs2 in the IPCs also rescues the *Fmr1^3^* phenotype, suggesting functional overlapping of the two genes. Our studies provide the first evidence that Tet can control the guidance of axons in the developing brain by modulating glutamatergic signaling and the function is mediated by its DNA-binding domain.

## Introduction

Ten-eleven translocation (TET) is one of the most important epigenetic and epi-transcriptomic regulators. TET can regulate DNA methylation in the genome as well as RNA hydroxymethylation through its catalytic activity ^1–3^. TET is an essential gene for animal development. Loss of all three TET genes in mice leads to embryonic lethality ^4^. *Drosophila* has one highly conserved Tet homolog and the *Tet^null^* mutant dies at the late pupal stage ^5^. Mammalian TETs are expressed in various tissues while *Drosophila* Tet is highly expressed in neuronal tissues including the brain ^5, 6^. Recently, TET mutations in humans have been linked to neurodevelopmental disorders (ND) associated with intellectual disability and autism spectrum disorders ^7–10^. One key process during brain development is axon guidance and the misregulation of axon guidance can lead to neurodevelopmental disorders ^11^. To test whether Tet functions in axon guidance, we used a well-studied axonal structure in the *Drosophila* brain, the mushroom body (MB).

The insect MB is the analogous structure of the human hippocampus ^12^. In *Drosophila*, the MB is responsible for learning and memory, circadian rhythms, sleep, and courtship ^13, 14^. In each brain hemisphere, the MB structure develops from four embryonic MB neuroblasts ^15^. During the development, these MB neuroblasts sequentially generate different classes of MB neurons whose axons bundle into γ, α′/β′, and α/β lobes ^15^. These MB lobes can be divided into 15 compartments which are connected to specific MB output neurons or dopaminergic neurons as part of the brain circuit to modulate a variety of brain functions ^13^. Thus, MB axon guidance is highly regulated to ensure proper brain circuitry.

Glutamatergic signaling is the major excitatory pathway in the brain and has key roles in neuronal plasticity, development, learning and memory ^16–18^. The NMDA-type glutamate receptors (NMDARs) are present in axons and growth cones of young hippocampal neurons ^19^. Mis-regulation of the metabotropic glutamate receptor (mGluR) signaling causes neuronal defects ^20^. Glutamate levels can affect the growth rate and branching of dopaminergic neurons *in vitro* ^21^. In addition, the responsiveness of growth cones to repellent molecules such as slit-2, semaphoring 3A, and semaphoring 3C is dependent on the levels of glutamate in the growth cone culture medium ^22^. However, there is limited *in vivo* evidence on whether and how glutamatergic signaling can control axon guidance.

Fragile X Messenger Ribonucleoprotein 1 (FMR1) is a human gene encoding the Fragile X Messenger Ribonucleoprotein (FMRP) which is important for regulating synaptic plasticity, protein synthesis, and dendritic morphology ^23^. In patients with Fragile X syndrome (FXS), the most common form of inherited intellectual and developmental disability, the gene generally is silenced by CGG trinucleotide expansion ^23^. FMR1 is conserved in *Drosophila* and mutants in the *Fmr1* homolog have been used to study the requirement of the gene and to identify potential therapeutic treatments for FXS ^24, 25^. The *Drosophila Fmr1* mutants have been shown to have defects in axon guidance and to affect neurological functions such as learning and memory as well as courtship behavior ^20, 26^.

In this study, we investigate a new function of *Tet* in brain development. We found that Tet can control MB axon guidance by regulating glutamine synthetase 2 (Gs2), a key enzyme in glutamatergic signaling. Tet DNA-binding domain is mainly responsible for the function. The regulation occurs during early brain development and mediated by the insulin-producing cells (IPCs). Modulating glutamatergic signaling by treating *Tet* mutant with mGluR inhibitor MPEP or glutamate can rescue or enhance the axon guidance defects in the mutant. Interestingly, *Tet* and *Fmr1* mutants exhibit similar phenotype and Gs2 reduction in the brain; and expressing Gs2 in the IPCs can rescue the phenotype in both mutants, suggesting a functional interaction of the two genes.

## Results

### Tet controls axon guidance in the brain via its DNA-binding domain

In the *Drosophila* brain, MB lobes are bundles of axons originating from MB neurons. In wild-type brains, axons of these lobes are separated at the brain midline, e.g., the β lobe in the left-brain hemisphere never touches the β lobe on the right side (**Figure 1A**). To test whether Tet is required for MB axon guidance, we knocked down its expression using RNAi under the control of 201Y-GAL4 driver. This driver is known to express in MB α/β core and γ neurons ^27^ and in midline cells (this study, **Figure 4A**). The driver was previously recombined with UAS-CD8::GFP to visualize its expression in MB cell bodies and axons ^27^. Tet knockdown using 201Y-GAL4 causes MB β axons to cross the midline and fuse with the β axonal bundle from the opposite brain hemisphere causing β lobe fusion (compare **Figure 1C, E, G** with **Figure 1B, D, F)**. 71% of Tet RNAi flies show β lobe fusion while none of the RNAi control flies showed fusion (**Figure 1H**). Tet RNAi did not affect γ or α lobes (**Figure S1A and B, Figure 1C-G**). Thus, Tet is required for the guidance of MB β axons.

**Figure 1.**
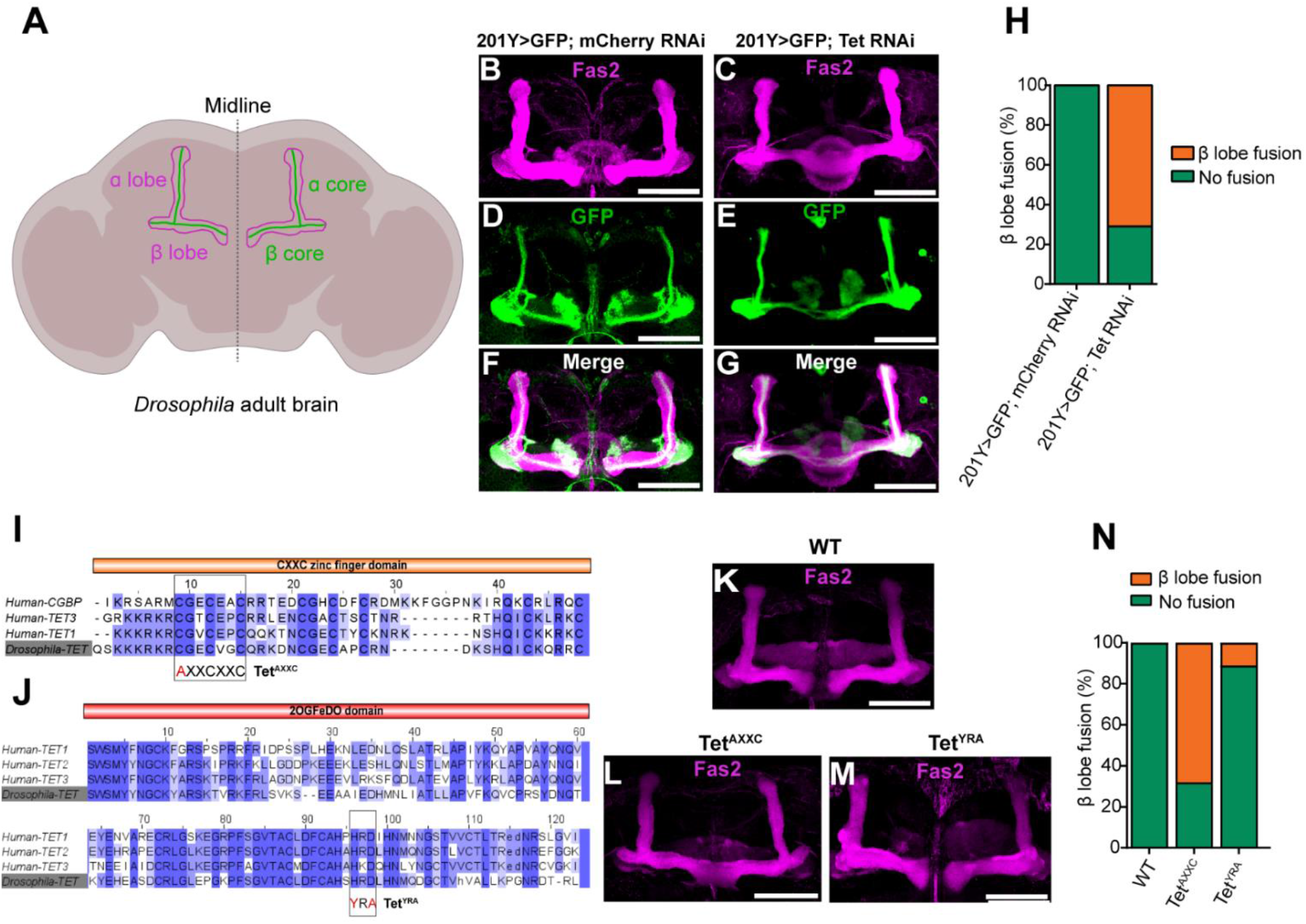
Tet and its DNA-binding domain are required for proper MB β axon guidance. (**A**) Schematic representation of the adult MB α/β lobe structure and the brain midline, the α/β lobes are shown in anti-magenta, α/β core axons are shown in green. (**B-G**) Tet knockdown (KD) using the 201Y- GAL4 MB driver, which also controls the expression of membrane-tethered CD8::GFP. α/β lobes were stained with anti-Fas2 antibody (B, C, magenta), α/β cores were stained with anti-GFP antibody (D, E, green), F and G are merged images of B and D or C and E. (**H**) Percentage of β lobe fusion phenotype obtained from RNAi control (n=29) or Tet KD (n=31) in (B-G). (**I**) Sequence homology and amino acid conservation of Tet and CGBP CXXC zinc finger DNA-binding domains. (**J**) Sequence homology and amino acid conservation of human and *Drosophila* Tet 2OGFeDO catalytic domains. (**K-M**) MB α/β lobes stained with the anti-Fas2 antibody of Tet domain mutants: DNA-binding domain C to A mutation (*Tet^AXXC^*), or catalytic domain HRD to YRA mutation (*Tet^YRA^*). (**N**) Quantification of K-M, percentage of normal or β fusion phenotypes, wild-type *WT* (n=30), *Tet^AXXC^*(n=37), and *Tet^YRA^* (n=35). Scale bar: 50 µm.

Tet has two domains, CXXC zinc finger and 2OGFeDO which are respectively responsible for DNA- binding and catalytic activities. The mammalian TET DNA-binding domain can bind to DNA to modulate epigenetic marks or to recruit other epigenetic factors to regulate gene expression ^28–30^. The catalytic domain is responsible for converting 5-methylcytosine (5mC) to 5-hydroxymethylcytosine (5hmC) in DNA and RNA ^1, 2^. To investigate the functions of these domains, we employed a CRISPR/Cas9 and homologous directed repair (HDR) approach to generate targeted point mutations ^31–33^. A previous study had shown that mutating one cysteine in the highly conserved zinc finger domain of the CGBP (CpG- binding protein) protein to alanine completely abolished its ability to bind DNA (**Figure 1I**) ^34^. We generated a mutant fly line in which the same conserved cysteine within Tet was substituted by alanine and called the mutation *Tet^AXXC^* (**Figure 1I**). Using the same CRISPR/Cas9 and HDR approach, we also generated a mutant in which the conserved histidine and aspartic acid in the active site of the catalytic domain were substituted by tyrosine and alanine and called the mutation *Tet^YRA^* (**Figure 1J**). This change in amino acids was previously shown to abolish the catalytic function of vertebrate Tet ^1^. Both these homozygous mutant fly lines show only partial lethality, unlike homozygous *Tet^null^* flies that are completely lethal and die at late pupal stage (**Figure S1D and E**). We analyzed the changes in MB structure in both mutant lines. By staining the adult brains with anti-Fas2 antibody to visualize the α/β lobes, we found that homozygous *Tet^AXXC^* showed the β lobe fusion phenotype in 68%, while, surprisingly, in homozygous *Tet^YRA^* the fusion was seen in just 11%, and wild type had 0% β lobe fusion (**Figure 1K-M and N**). Thus, Tet DNA-binding domain is primarily required for MB β axon guidance.

### Tet is required in early brain development to prevent β axons crossing the brain midline

MB development starts in early embryogenesis and originates from four neuroblasts (**Figure 2A**). To determine when during development Tet is required for β axon guidance, we combined the 201Y-GAL4 with temperature sensitive GAL80 (GAL80^ts^) for a temperature shift experiment allowing stage specific RNAi induction. We selectively induced Tet RNAi from white prepupal stage (WPP) to adult, WPP to 48h after puparium formation (APF), or embryo to WPP (**Figure 2B**). Tet RNAi during embryogenesis and larval development does not cause β axon midline crossing (embryo to WPP) but Tet RNAi during pupation (WPP to adult, WPP to 48h APF) resulted in β fusion observed in the adult brain, similar to what is seen when Tet is depleted throughout development (embryo to adult) (**Figure 2C**). Tet is therefore specifically required in early pupae for normal β axon guidance. To confirm the temperature shift result, we also dissected and stained *Tet^AXXC^* brains with anti-Fas2 antibody at four time points between WPP and 48h APF. We first observed β axons crossing the midline in the *Tet^AXXC^* brain 24h APF (**Figure 2D**). Thus, Tet is required during the first 24h APF to prevent β axons from crossing the brain midline.

**Figure 2.**
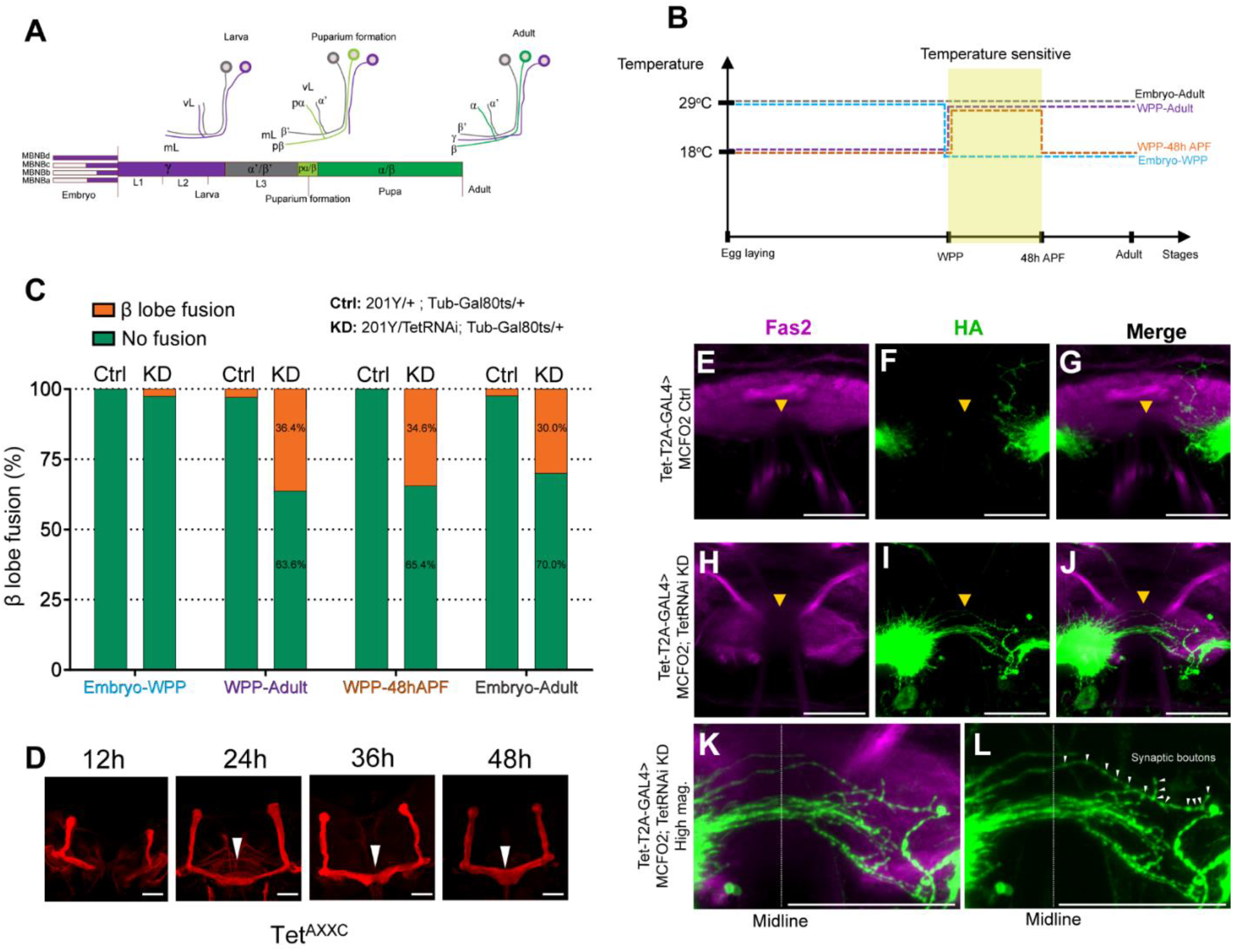
Tet is required in the first 24h APF to prevent β axons from crossing over the brain midline. (**A**) The progression of MB axon structural development from four embryonic MB neuroblasts (MBNBs) is simplified and depicted following the Kunz et.al. 2012 study ^15^. (**B**) Temperature shift (TS) scheme to knockdown Tet using the GAL4/GAL80^ts^ system at four developmental stages: embryo to adult, WPP to adult, WPP to 48h APF, embryo to WPP. WPP: white prepupa, APF: after puparium formation. (**C**) Percentage of brains with β lobe midline crossing phenotype (orange) in TS Tet KD (201Y> TetRNAi; tub-GAL80^ts^, KD) and control (201Y> tub-GAL80ts, Ctrl). Tet KD from embryo to WPP does not affect the growth of β axons, but Tet KD from embryo to adult, WPP to adult, or WPP to 48h APF resulted in β axon midline crossing. The temperature-sensitive period is indicated in yellow in (**B**). (**D**) *Tet^AXXC^* pupal brains from different developmental stages (12h, 24h, 36h, 48h APF) were stained with anti-Fas2 antibody. β axon midline crossing occurs from 24h APF onward. (**E-M**) Stochastic labeling of MB β axons in the developing pupal brain (21h APF), Tet KD driven by Tet-T2A-GAL4 (**H-J**, Tet-GAL4> MCFO2, TetRNAi) or control (**E-G**, Tet-GAL4> MCFO2) brains. Brains were stained with anti-HA antibody to label axons (green) and anti-Fas2 antibody (magenta) to label MB and ellipsoid bodies to identify the brain midline. Tet KD in the developing brain leads to growing axons crossing the midline (**I and J**) not observed in the control (**F and G**). (**K**) and (**L**) are high magnifications of (**J**) and (**I**). Tracing of one axon in (**K**) showing the portion of the axon that has crossed the midline. Note the synaptic boutons distributed along it (**L**, arrowhead). Scale bars: 25 µm.

Anti-Fas2 staining labels MB axon bundles; however, it cannot reveal how single axons grow. Thus, we employed a stochastic labeling method in which a membrane tethered molecular marker called smGdP (spaghetti monster Green darkened Protein) was put under the control of the GAL4/UAS and heat shock FLP/FRT inducible systems ^35^. This method has been used to label single neurons, axons, and small dendrites at high resolution ^35^. To label growing axons, we need a GAL4 driver that is expressed in neurons that have active axon growth. Tet-T2A-GAL4, which is a GAL4 driver expressed under the control of the Tet promoter ^36, 37^, expresses in such neurons. We expressed smGdP under Tet-T2A-GAL4 control, heat shocked at WPP, and stained brains at 21h APF with anti-HA antibody. The result showed that Tet-T2A-GAL4 drove expression in a subset of MB neurons, their axons (**Figure S2B**) and importantly, growing axons (**Figure S2C-E)** including β axons projecting their growth cone toward the brain midline (**Figure S2B and D**). In addition, inducing Tet RNAi by Tet-T2A-GAL4 produced the same β lobe midline crossing phenotype as seen with the 201Y-GAL4 driver (**Figure S2F and G**). Thus, we can use Tet-T2A-GAL4 driver for simultaneously inducing Tet RNAi and labeling growing β axons to see how these axons behave under wild type versus Tet RNAi conditions. We found that in Tet RNAi at 21h APF, some axons at the tip of the left β lobe are protruding and extending their projections over the midline toward the right β lobe (**Figure 2H-J**). This phenomenon does not occur in the control (**Figure 2E-G**). In this experiment we look at an early time in prospective adult axon outgrowth and therefore the very first axons crossing the midline. These axons will serve as a guide for other axons to cross, resulting in the complete fusion we observed in the adult brain (**Figure 1**). In addition to the Tet RNAi axon outgrowth defect, we also noticed that there are small swellings located along the portion of the axon that crossed the midline (**Figure 2K and L**, arrowheads). These swellings are the signs of synaptic boutons indicating that the midline crossing axons, normally restrained to the left-brain hemisphere, likely form synapses with neurons from the right brain hemisphere and potentially affect the MB circuit. Thus, Tet is required to keep β axons confined to one of the two brain hemispheres to prevent any misconnection of β axons between the two sides of the brain.

### Tet regulates axon guidance by controlling the levels of glutamine synthetase 2, a key enzyme in glutamatergic signaling

To investigate how Tet controls MB axon guidance, 24h pupal brains of *wild-type* and *Tet^AXXC^* were dissected and subjected to RNA sequencing. The result showed that in the *Tet^AXXC^*brains, 1503 genes are up-regulated and 29 genes down-regulated compared to wild type (**Figure 3A and Supplemental data 1 and 2**). Most of the up-regulated genes are expressed at low levels and GO term analysis showed that these genes functioned in sperm development and motility (**Figure S3A and B**). But interestingly, the two most significantly down-regulated genes, *Glutamine synthetase 2* (*Gs2*) and *GABA transporter* (*Gat*) encode for the key enzymes of glutamatergic signaling and the key transporter of GABAergic signaling (**Figure 3A and B**). Gs2 can convert glutamate to glutamine and Gat can uptake GABA (**Figure 3B**) ^38–40^, suggesting a new and interesting regulation of *Tet* in glutamatergic and GABAergic signaling that potentially affects axon guidance.

**Figure 3.**
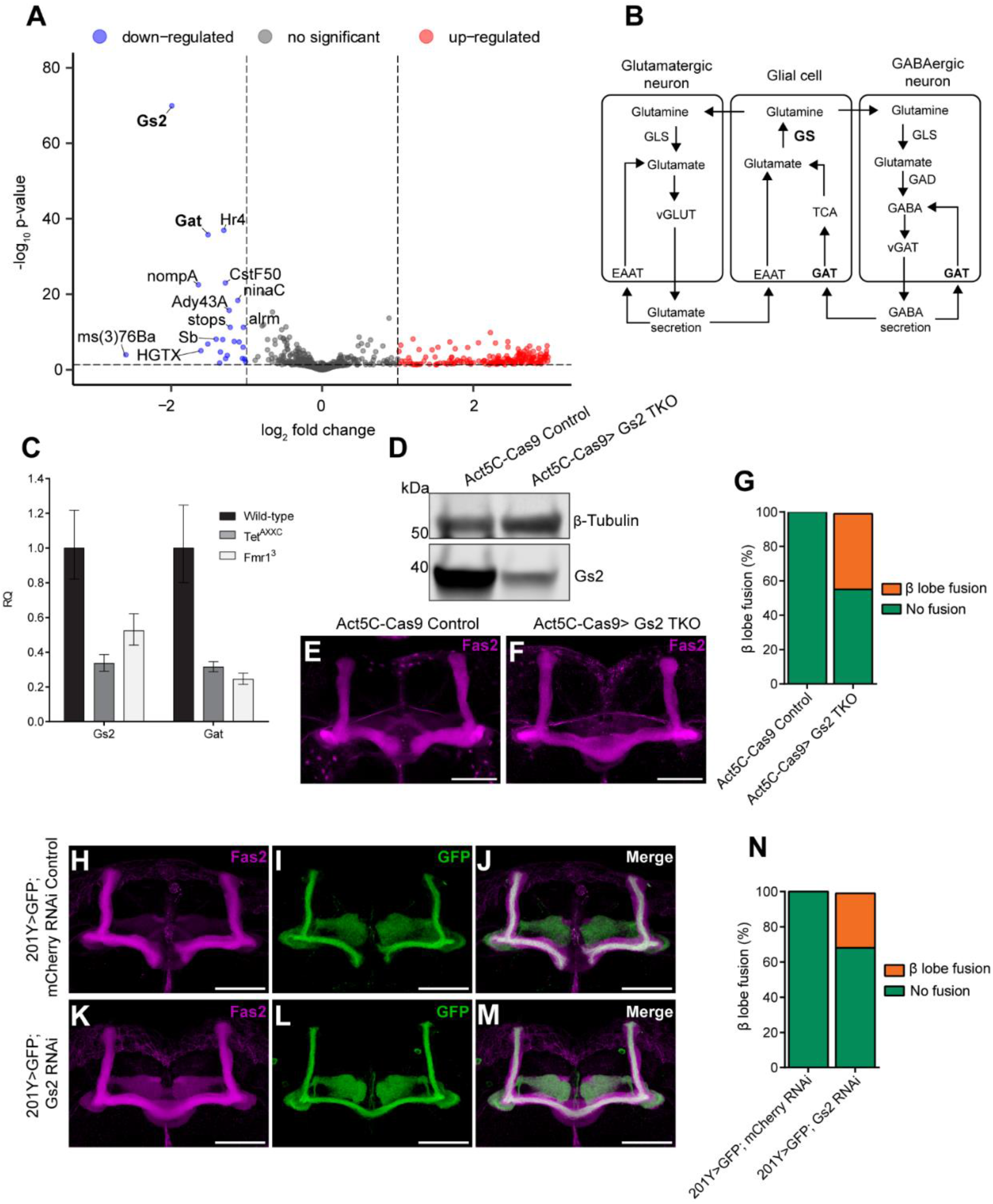
Tet regulates β axon guidance by controlling glutamate synthetase 2 levels in the brain. (**A**) Volcano plot of RNA sequencing results, showing differentially expressed RNAs in *Tet^AXXC^* and *wild- type* brains. Significant differences are defined by p-value < 0.01 and fold change > 2. (**B**) Representation of the glutamatergic and GABAergic pathways showing that Gs2 and Gat are the main enzyme and transporter, respectively, of the pathway. (**C**) Comparison of Gs2 and Gat transcript levels in *wild-type* or *Tet^AXXC^*, and *Fmr1^3^* mutant brains assessed by RT-qPCR. RQ: relative quantification, error bars represent standard deviation (SD). (**D**) Western blot showing Gs2 protein levels in the brains of control (Act5C- Cas9) or CRISPR/Cas9 mutated Gs2 (Act5C-Cas9> Gs2 TKO) blotted with anti-Gs2 antibody. (**E-F**) MB α/β lobes stained with anti-Fas2 antibody showing Gs2 mutated brain exhibiting MB β lobe midline crossing (**F**, Act5C-Cas9> Gs2 TKO) not observed in control brains (**E**, Act5C-Cas9 Control). (**G**) Quantification of phenotype seen in (E-F). (**H-M**) Gs2 RNAi KD driven by 201Y-GAL4 (K-M) or control (H-J). All brains express 201Y-GAL4 driven CD8::GFP. Brains were stained with anti-Fas2 and anti-GFP antibodies to visualize whole α/β lobes (Fas2, magenta) or core α/β (GFP, green). Gs2 KD using MB driver 201Y-GAL4 recapitulates the *Tet^AXXC^*midline crossing phenotype not observed in control (K-M). (**N**) Quantification of midline crossing phenotype in (H-M). Scale bars: 50 µm.

To confirm the RNA sequencing results, we performed RT-qPCR to test the RNA levels of *Gs2* and *Gat* in *wild type* and *Tet^AXXC^* brains. The result confirmed that both genes are down-regulated in *Tet^AXXC^* compared to *wild type* (**Figure 3C**). In addition, because it had been shown that *Fmr1* mutants also exhibit the β lobe fusion phenotype ^20, 41^, we also dissected 24h pupal brains of *Fmr1^3^*mutant ^24^ to test the expression of these same two genes. Interestingly, *Gs2* and *Gat* are also down-regulated in *Fmr1^3^* (**Figure 3C**).

Glutamate can control the growth rate of dopaminergic axons ^21^ and can also reduce the responsiveness of growth cones to repellent cues ^22^. Further, GABA has been shown to control axonal growth ^42, 43^. Thus, the down-regulation of *Gs2* and *Gat* may be responsible for the MB β axon midline crossing phenotype. To test this hypothesis, we expressed *Gs2* and *Gat* RNAis under the control of the 201Y-GAL4 driver. *Gs2* RNAi produced the β axon midline crossing phenotype similarly to what is observed in the *Tet^AXXC^* mutant (**Figure 3H-M and N**) indicating Tet regulates β axon guidance via Gs2. Due to lethality that occurred in *Gat* RNAi animals, we were not able to determine the requirement of *Gat*; thus, we focused on *Gs2* for further experiments.

To confirm the *Gs2* RNAi phenotype, we employed CRISPR/Cas9-mediated mutagenesis to inactivate the *Gs2* gene ubiquitously. The mutagenesis is mediated by the combination of a ubiquitous Cas9 expression driver Act5C-Cas9 and a ubiquitous Gs2 sgRNA expression stock. The Gs2 mutagenized brains showed a significant reduction of Gs2 protein compared to control brains (**Figure 3D**). Importantly, 45% of the *Gs2* mutagenized brains showed β axons crossing the midline while no control brains showed this phenotype (**Figure 3E, F, and G**). These results validate Tet function in controlling MB β axon guidance by regulating the level of Gs2 in the brain.

### Tet acts in insulin-producing cells (IPCs) to regulate axon guidance

Expressing *Tet* or *Gs2* RNAi under 201Y-GAL4 driver results in the β lobe midline crossing phenotype. However, exactly which cells in the brain express Gs2 to control MB organization is not known. At 24h APF, 201Y-GAL4 expresses in both MB neurons and a group of cells at the midline of the brain named midline cells (MLCs) (**Figure 4A**). To examine if the MB neurons or MLCs express Gs2, brains were stained with anti-Gs2 antibody ^44^. In brains expressing CD8::GFP driven by 201Y-GAL4, the Gs2 antibody does not stain the MB neurons (**Figure 4B-D**) but strongly stains the MLCs (**Figure 4E-G**, asterisks). Similar to these MLCs, another MB driver, OK107-GAL4, also expresses in the median neurosecretory cells which mostly identified as insulin-producing cells (IPCs) ^45^. In addition, Cao et al., have shown that Gs2 is highly enriched in IPCs ^46^. Thus, we suspect that the MLCs identified by 201Y- GAL4 are IPCs and hypothesize that Tet regulates Gs2 in these cells to control MB β axon guidance. To test this hypothesis, we drove *Tet* or *Gs2* RNAis using a Dilp2-GAL4 driver, which is expressed specifically in IPCs. Expressing *Tet* or *Gs2* RNAis in the IPCs produced an MB β axon crossing phenotype similar to that observed in *Tet^AXXC^* (compare **Figure 4I, J, and K** to control **Figure 4H and K**). This result indicates that Tet and Gs2 are required in the IPCs to regulate MB axon guidance.

### *Tet* and *Fmr1* mutant phenotypes can be rescued by expressing Gs2 in IPCs

To validate that Tet and Gs2 act in IPCs to regulate β axon guidance, we generated a Gs2 transgene and expressed it in the IPCs to test if we can rescue the *Tet^AXXC^* mutant phenotype. For these rescue experiments, *Tet^AXXC^*mutants that expressed Gs2 in the IPCs (*Dilp2-GAL4/UAS-Gs2; Tet^AXXC^/ Tet^AXXC^*) (**Figure 4N**) were compared to the mutant control, *Tet^AXXC^* mutants that did not express Gs2 (*Dilp2- GAL4/+; Tet^AXXC^/ Tet^AXXC^*) (**Figure 4M**) and the wild-type control (*Dilp2-GAL4/+*) (**Figure 4L**). Since *Tet* and *Fmr1* mutants exhibit similar phenotypes as down regulation of Gs2, the same experimental design was applied to rescue *Fmr1^3^*mutants (**Figure 4L-N**). Results showed that while Dilp2-GAL4 alone did not affect the β axon guidance, expressing Gs2 in the IPCs of *Tet^AXXC^* brains can significantly reduce the *Tet^AXXC^* phenotype (26% MB fusion) compared to *Tet^AXXC^*flies without Gs2 expression (47% MB fusion, **Figure 4O**). Furthermore, when Gs2 is expressed in the IPCs of the *Fmr1^3^* mutant brains, the structural defect was also reduced (21% MB fusion) compared to the homozygous mutant *Fmr1^3^* brains (54% MB fusion, **Figure 4P**). Taken together, these results demonstrate that Tet and Gs2 act in the IPCs to regulate MB β axon guidance. The results also suggest an important interaction of Tet and Fmr1 in regulating axon guidance via Gs2.

**Figure 4.**
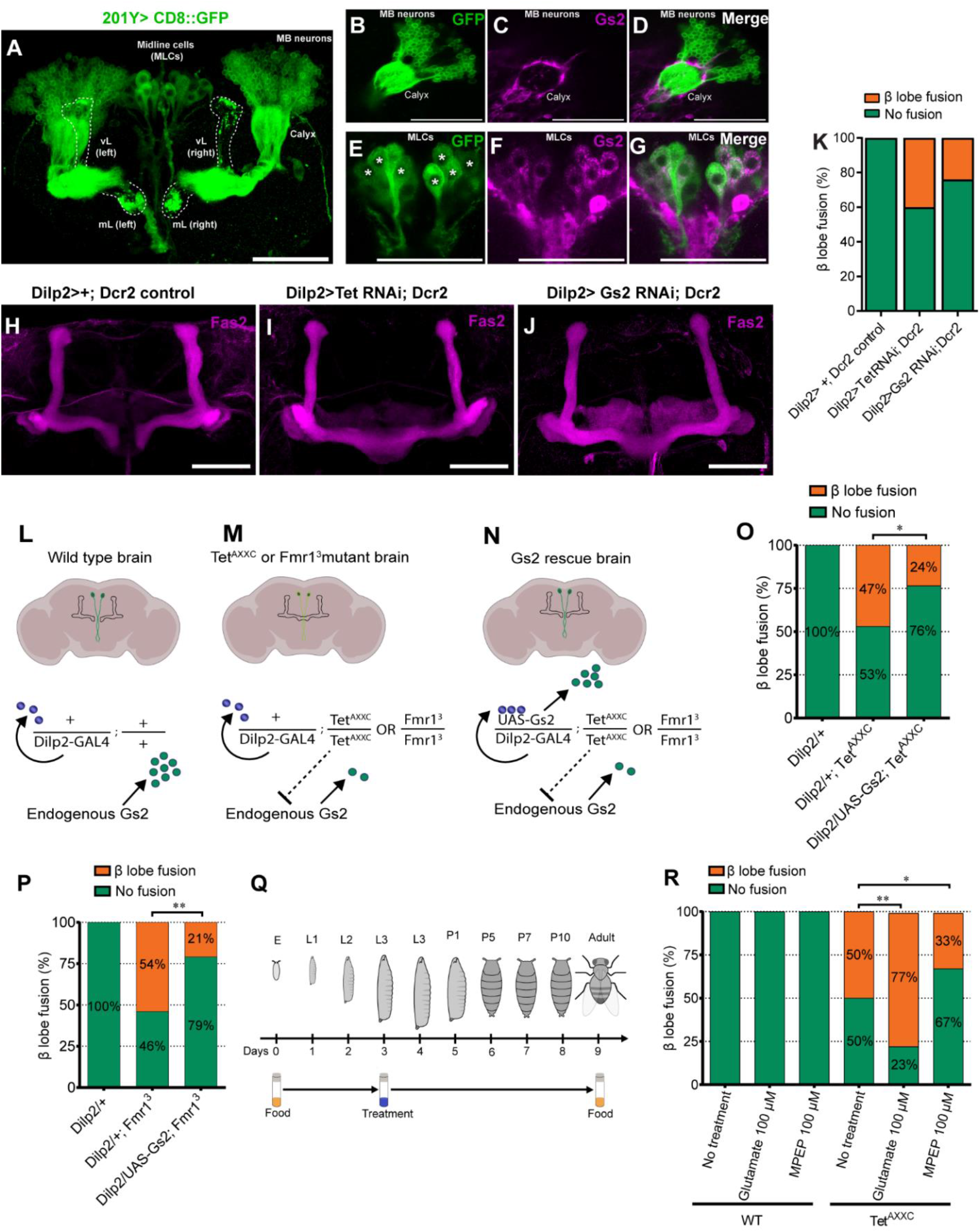
Tet regulates β axon guidance via insulin-producing cells (IPCs) and the *Tet* and *Fmr1* mutant phenotypes can be rescued by over-expressing Gs2 in the IPCs or MPEP treatment. (**A**) 201Y-GAL4 driven membrane-tagged CD8::GFP expression pattern in 24h APF brain, the driver expresses in MB neurons and midline cells (MLCs). vL: vertical lobe, mL: medial lobe. (**B-G**) 201Y-GAL4 driven CD8::GFP cells (green) and Gs2 expressing cells (magenta, anti-Gs2 staining). Gs2 is not expressed in MB neurons (B-D) but instead is expressed in MLCs (E-G, asterisks marks midline cells expressing Gs2). (**H-J**) MB α/β lobes stained with anti-Fas2 antibody in the Dilp2-GAL4 driver control (H), Dilp2-GAL4 driven Tet RNAi (I), or Dilp2-GAL4 driven Gs2 RNAi (J). (**K**) Quantification of β fusion brain in (H-J). (**L-N**) Schematic representation of the experimental design for rescuing *Tet* or *Fmr1* mutant phenotypes by expressing Gs2 under the control of Dilp2-GAL4. (**O**) Percentage of brains showing MB β axon midline crossing from wild type, *Tet^AXXC^* mutants, and Gs2 rescued *Tet^AXXC^* mutants. (**P**) Percentage of brains from wild type, *Fmr1^3^* mutants, and Gs2 rescued *Fmr1^3^*mutants showing MB β axon midline crossing. (**Q**) MPEP and glutamate treatment scheme for *wild-type* and *Tet^AXXC^* mutant flies. Third instar larvae were reared on food containing MPEP or glutamate and changed to normal food after eclosion. (**R**) Percentage of *wild type* or *Tet^AXXC^* mutant brains showing MB β axon midline crossing after MPEP or glutamate treatment. Chi-square test with * p < 0.05, ** p < 0.01. Scale bar: 50 µm.

### Axon guidance defects in *Tet* mutants are modulated by glutamatergic signaling

Gs2 is a key enzyme in glutamatergic signaling. Our results showed that Gs2 is regulated by Tet to control β axon guidance. In *Tet* mutant brains, the reduction of Gs2 may lead to the accumulation of glutamate and upregulation of glutamatergic signaling. Excess amounts of glutamatergic signaling have been proposed as the main cause of Fragile X syndrome ^47^. In fact, treating *Fmr1* mutants with metabotropic glutamate receptor (mGluR) antagonist MPEP can suppress the *Fmr1* mutant phenotypes ^20^ while treating the mutants with glutamate can enhance the phenotypes ^48^. To validate the function of Tet in modulating glutamatergic signaling, we examined if mGluR antagonist MPEP and glutamate can affect the *Tet* mutant phenotype. *Wild-type* and *Tet^AXXC^*third instar larvae were transferred to food supplemented with MPEP or glutamate as treatments, or water as control (**Figure 4Q**). The treatment had no negative effect on viability (**Figure S4**). Brains from 5 to 10-day-old flies were dissected and stained with anti-Fas2 antibody and the percentage of brains showing MB β midline crossing was quantified. The result showed that treating *Tet^AXXC^*mutants with glutamate increased the number of flies with the β axon crossing midline phenotype by 27% while treating with MPEP showed 17% reduction of the phenotype compared to the non-treated control (**Figure 4R**). Neither of the treatments affected *wild-type* brains (**Figure 4R**). Thus, modulating glutamatergic signaling by glutamate or the mGluR antagonist MPEP can enhance or suppress the β axon guidance defects in *Tet^AXXC^* mutant. This result confirms the function of Tet in regulating glutamatergic signaling to control axon guidance in the brain.

## Discussion

### Tet controls axon guidance during early brain development via its DNA-binding domain

TET is essential for neurodevelopment ^49^; but how Tet regulates axon guidance in the developing brain is understudied. Some examples are TET3 is upregulated and required for axon regeneration in adult dorsal root ganglion (DRG) neurons after injury ^50^ and Tet involves in commissure axon patterning and glial homeostasis in *Drosophila* ^51, 52^. However, the requirement of Tet and its DNA-binding domain in regulating axon guidance in the developing brain is not studied. Our results establish that Tet regulates the guidance of β axons in the MB, a prominent structure in the brain that controls different neurological functions in the brain ^13, 14^, and the regulation is dependent on its DNA-binding domain.

We found that Tet is required during early pupal brain development for normal MB β axon guidance. This is also the time point when the β axons grow to form the mature MB β lobes. This observation shows that the protein is required at a specific stage during brain development to guide new axons to the right destination. Our results also show that the requirement for Tet function in the MB β axon guidance is limited to the first 48 hours of pupal brain development. Careful inspection of the Tet RNAi axons that cross the midline showed that these axons formed small swellings, a sign of synaptic boutons, in the “wrong” brain hemisphere suggesting that the circuits of the hemispheres are misconnected. These observations suggest that treatment for neurodevelopmental disorders that are due to axon guidance problems and the ensuing misconnection of synapses, needs to be applied at early stages of nervous system development.

### Tet regulates glutamatergic signaling

Our transcriptomic studies demonstrated that *Gs2* and *Gat* are regulated by *Tet*. Gs2 is a key enzyme in regulating glutamate levels in glutamatergic signaling and Gat is the neurotransporter that controls GABA levels in the GABAergic signaling suggesting that Tet is required in both pathways. The function of Gat in controlling axon development needs to be further elucidated due to the lethal phenotype in Gat knockdown flies. Inducing Gs2 RNAi by the 201Y-GAL4 driver resulted in axon defects indicating that MB axon guidance is regulated by glutamatergic signaling via Gs2. In addition, treating *Tet^AXXC^*mutants with MPEP, a metabotropic glutamate receptor (mGluR) antagonist ^20^, can rescue the axon guidance defects while glutamate treatment enhances the *Tet^AXXC^* phenotype. These data demonstrate a new function of Tet in regulating glutamatergic signaling in the brain and support the notion that the pathway is mis-regulated in Tet mutant brains. Glutamatergic signaling is a major excitatory signal in the human brain and the pathway is also known to act in neural developmental events such as proliferation, differentiation, migration, and synaptogenesis ^18^. Interestingly, elevated levels of glutamatergic signaling have also been proposed as the underlying mechanism of the neurodevelopmental disorder in Fragile X syndrome ^47^. Based on the conserved functional domains of Tet in humans and flies, the mis-regulation of the glutamatergic signaling we observed in *Drosophila* brain during early development may also occur in developing human brains harboring Tet mutations leading to neurodevelopmental disorders.

We have previously mapped Tet-DNA binding sites in the genome by ChIP-seq and mapped Tet- mediated 5hmrC RNA modification throughout the transcriptome by hMeRIP-seq ^53^. There is no Tet binding detected by ChIP-seq on the gene body or promoter region of *Gs2* and *Gat* (**Figure S3C-E**). In addition, the Gs2 and Gat mRNAs do not carry the Tet-dependent 5hmrC modification ^53^. These observations suggest that Tet regulates *Gs2* and *Gat* indirectly. Since *Gs2* and *Gat* are also downregulated in the *Fmr1^3^*mutant, *Tet* and *Fmr1* may function together to control the mRNA levels of these two genes. Fmr1 was previously found in the regulatory protein complex Not1/CCR4 ^54^, which is known to control all steps of gene expression from transcription in the nucleus to mRNA degradation in the cytoplasm depending on the specific associated proteins that interact with the complex ^55^. We also have previously also identified Fmr1 in a complex with Zinc finger protein RP-8 (Zfrp8), which functions in the formation of mRNA ribonucleoprotein (mRNP) complexes ^56^. Interestingly, Zfrp8 was co-immunoprecipitated with Tet (**Figure S3F**). Thus, Tet and Fmr1 may both coordinate with the Not1/CCR4 or other mRNP complexes to regulate the expression levels of *Gs2* and *Gat*.

### Tet and Gs2 are required in the insulin-producing cells (IPCs) to regulate axon guidance

IPCs in *Drosophila* are a group of neurons located at the dorsal midline of the central brain that produce and secrete insulin-like peptides into the hemolymph to regulate growth and metabolism ^57, 58^. Previous studies have shown that IPCs can also regulate neuronal functions ^59, 60^. For instance, overexpressing Fmr1 in IPCs could rescue the circadian defects observed in *Fmr1* mutants ^61^. But whether the IPCs can regulate MB axon guidance has not been elucidated. In our RNA-seq experiments, *Gs2* is the most significantly reduced transcript in the *Tet^AXXC^* mutant brain. We found that Gs2 is expressed in the IPCs but not in MB neurons. We observed the axon midline-crossing phenotype when we specifically knock down either Tet or Gs2 in IPCs. In addition, the axon midline-crossing phenotype was rescued when expressing Gs2 in the IPCs of *Tet* and *Fmr1* mutants. Hence, our results point to a previously unknown function of IPCs in regulating MB axon guidance via glutamatergic signaling in the developing brain.

Our result shows that Tet can control the levels of Gs2 and is required in the IPCs influence the guidance of the β axons. Gs2 is known expressed in glial cells where it is responsible for converting glutamate to glutamine to regulate the levels of the neurotransmitter glutamate in the brain ^62, 63^; however, it is also known to be highly enriched in IPCs ^46^. Thus, the reduction of Gs2 levels in *Tet^AXXC^* mutant may lead to the accumulation of glutamate and a deficiency of glutamine in the IPCs. However, the mechanism by which Tet and Gs2 regulate axon guidance via IPCs needs to be further elucidated. One possible mechanism is that, in Tet mutant brains, Gs2 is reduced in the IPCs leading to the accumulation of glutamate in the IPCs that is then secreted at the brain midline. The elevated levels of glutamate at the midline could cause MB β axons to cross over the midline. In suppost of this hypothetic mechanism, glutamate has been reported to reduce the responsiveness of axonal growth cones to repellent cues such as Slit-2 and Sema-3A/C ^22^ and also can enhance the growth rate and branching of dopaminergic axons ^21^. In addition, glutamate can be released by pancreatic cells, which are the insulin secreting cells in mammals, via excitatory amino acid transporters (EAAT) by uptake reversal ^64^. The uptake reversal activity of EAAT has also been reported in astrocyte and it depends on the relative intracellular and extracellular glutamate concentration ^65^. Further studies are needed to test the secretion of glutamate by IPCs, and to investigate whether the MB β axon crossing phenotype is due to elevated glutamate levels at the midline of *Tet^AXXC^* mutant brains.

### Potential interplay between Tet and Fmr1

Previously, MPEP was successfully used to suppress the axonal defect in *Fmr1^3^* mutants ^20^ while high glutamate concentration can enhance the phenotype ^48^. Our observation that MPEP and glutamate have similar effects on the *Tet^AXXC^* phenotype suggests that Tet and Fmr1 have overlapping functions. This conclusion is further strengthened by our finding that Gs2 is downregulated in the brain of both *Tet^AXXC^*and *Fmr1^3^* mutants, and that increasing Gs2 levels by over-expressing it from a transgene reduced the occurrence of the β lobe fusion phenotype in both *Tet^AXXC^* and *Fmr1^3^* mutants (**Figure 5L-P**). The rescue is partial, suggesting that the expression levels of Gs2 are essential or additional targets of Tet and Fmr1 also contribute to the phenotype. As Tet and Fmr1 can regulate a broad spectrum of genes, it is impressive that overexpression of only one gene, Gs2, encoding a key enzyme in the glutamatergic pathway, can rescue the axonal defect in about half of both *Tet* and *Fmr1* mutants. Dysfunction of glutamatergic signaling is involved in a variety of human diseases including Fragile X syndrome, autism spectrum disorder, schizophrenia, neurodegenerative diseases, and gliomas ^47, 66–68^. Our results suggest that glutamine synthetase is a potential target for early intervention. Recently, an increasing number of studies have shown that Tet mutations are associated with neurodevelopmental disorders in humans ^7–10^. The mechanism by which human TET mutations cause these disorders needs to be elucidated, but the highly conserved functional domain of human and *Drosophila* Tet suggests that the Tet function in axon guidance that we found may be conserved in humans. Thus, our study provides initial evidence that glutamatergic signaling is altered in the Tet mutant brains and the mutant may serve as a *Drosophila* model for drug screening and study of neurodevelopmental disorders caused by TET mutations.

## Materials and Methods

### Drosophila strains

Fly stocks were cultured on the standard food medium at 25 °C. The following stocks were obtained from Bloomington Drosophila Stock Center (BDSC): UAS-mCD8::GFP, 201Y-GAL4 driver (#64296) ^27^, enhancer trap Tet-T2A-GAL4-pA (#76666) ^37^, multicolor flip-out stock MCFO-2 (#64086) ^35^, Act5C-Cas9 (#54590), Gs2 sgRNA for Gs2 CRISPR/Cas9 mutagenesis (#76492), Gs2 RNAi (#92838) ^69^, Gat RNAi (#29422), mCherry RNAi (#35785), tubP-GAL80^ts^ (#7017), and UAS-Dcr2 (#24651). Tet RNAi lines were obtained from Vienna Drosophila Resource Center (VDRC, GD#14330 and KK#108964). Dilp2-GAL4 was originally obtained from Dr. Eric Rulifson (University of California, San Francisco) ^57^. Fmr1^3^ mutant was kindly provided by Dr. Thomas A. Jongens (University of Pennsylvania) ^24^.

### Conditional knockdown using temperature shift GAL80^ts^

The driver 201Y-GAL4 was crossed with *Tet RNAi (*GD#14330*); GAL80ts* or *mCherry RNAi; GAL80ts* to generate Tet knockdown GAL80^ts^ or mCherry RNAi control GAL80^ts^. The crosses and progenies were kept at 18°C or 29°C following the temperature shift scheme detailed in (**Figure 2B**). Five to ten days old adult brains were dissected and stained to calculate the percentage of β fusion phenotype on each condition.

### Generating targeted Tet^AXXC^ and Tet^YRA^ mutants using CRISPR/Cas9 and HDR

The *Tet^AXXC^* mutant was generated using the scarless gene editing approach which is mediated by CRISPR/Cas9 cut and homologous directed repair (HDR) of the designed DNA template harboring targeted mutations ^31–33^. Two gRNAs (gRNA1 and gRNA2) were designed to bind to both sides of the CXXC zinc finger domain flanking regions to guide the Cas9 cut. These gRNAs were synthesized (IDT, USA) and cloned to the pU6-BbsI-chiRNA expression vector (Addgene #45946)^33^. The two homology arms were generated by PCR from the nos-Cas9 stock (BDSC # 54591) genomic DNA. The PCR was done using gene-specific and mutated primers to generate one wild-type homology arm and one mutated homology arm with at least 1000 bp overlap with the flanking genomic DNA from the cut site (Fig. S7). The homology arms were cloned to the pScarlessHD-DsRed donor vector (Addgene # 64703, gift from Kate O’Connor-Giles). The mixture of these vectors was injected into the nos-Cas9 stock and screened for the DsRed marker. DsRed positive lines were genetically crossed to remove the non-Cas9 chromosome. Then, these lines were crossed to a piggyBac recombinase stock 3xP3-ECFP, alpha-tub- piggyBacK10/TM6C, Sb^1^ (BDSC # 32072) to excise the DsRed marker. The mutant stocks were validated by PCR and Sanger sequencing. The *Tet^YRA^* mutant was generated using the same method. The sequences for sgRNAs and primers to generate homology arms are listed in **Supplementary Table S1**.

### Simultaneously knockdown and visualizing growing axons using Tet-T2A-GAL4

The Tet enhancer trap line Tet-T2A-GAL4-pA ^37^ was used as Tet-T2A-GAL4 driver to drive Tet knockdown at Tet-expressing cells in the pupal brain and labeling cell body as well as axons of these cells. The knockdown can be accomplished because Tet-T2A-GAL4-pA was inserted between exons 5 and 6 but the RNAi stock #110549 (VDRC, construct KK#108964) was bound to exon 8 (**Figure S2A**). The poly(A) (pA) signal will trigger the transcriptional termination to ensure the RNAi will not bind and degrade the transcript containing the GAL4 coding sequence. To inducible label axons with high resolution, we employed the MCFO2 stock which can express membrane tethered smGdPs (spaghetti monster Green darkened Proteins) under the control of both GAL4/UAS and heat shock FLP/FRT inducible systems ^35^. smGdP is a darkened chromophore Green Protein fused with multiple copies of epitope tag (e.g., HA, V5, OLLAS, FLAG, and Myc) and a myristoylation for better visualization of small axons and dendrites ^70^. The Tet RNAi stock (KK#108964) or mCherry RNAi were crossed to the flip-out stock MCFO-2 to generate *UAS-Tet RNAi; MCFO-2* or *UAS-mCherry RNAi; MCFO-2* stocks. Then, Tet- GAL4 was crossed to the *UAS-Tet RNAi; MCFO-2* for Tet knockdown and labeling axons of Tet expressing neurons, or *UAS-mCherry RNAi; MCFO-2* for the control. The crosses were kept at 25°C to produce white pupae. Then, white pupae were collected and placed on the wall of the *Drosophila* glass vials. The vials were heat shocked for 3 minutes at 37°C in a water bath, then transferred to a 25°C incubator and grown for 21 hours. Then, pupal brains were dissected and stained for visualizing the growing MB axons.

### Brain dissection, immunostaining and imaging

Brain dissection and immunostaining were performed as previously described ^71^ with the following modifications. Adult brains were dissected in cold 1X phosphate-buffered saline (PBS) and fixed in 4% formaldehyde at 25°C for 20 min on a slow rotator (7 rpm). Pupa brains were dissected in cold 1X PBS, then immediately transferred to a cold fixation solution containing 4% formaldehyde diluted in Schneider’s *Drosophila* Medium (Gibco # 21720024) kept on ice. The pupa brains can be kept in the cold fixation solution for a maximum of one hour. Then these pupa brains were fixed at 25°C for 45 min on a slow rotator (7 rpm). After fixing, brains were quickly washed with 0.3% PBS-T (0.3% Triton-X100 in 1X PBS) one time, then three washes with 0.3% PBS-T, 20 min each. The brains were blocked in blocking solution (0.15% PBS-T containing 10% normal goat serum) at 25 °C for 20 min. These brains were then incubated with the following primary antibodies diluted in blocking solution: mouse anti-Fasciclin II (Fas2, DSHB #1D4, deposited by Corey Goodman, 1:50), rabbit anti-GFP (Invitrogen #A-11122, 1:1000), mouse anti- glutamine synthetase (GS) antibody, clone GS-6 (Millipore-Sigma #MAB302, 1:1000) at 4 °C overnight or rabbit anti-HA (Invitrogen #MA5-27915, 1:1000) at 4°C over two nights. After primary antibody incubation, the brains were washed with 0.3% PBS-T, 3 times for 20 minutes each. Then, brains were incubated with secondary antibodies conjugated with Cy3 (Jackson ImmunoResearch #115-165-003, 1:400), Cy5 (Jackson ImmunoResearch #711-175-152, 1:400), or Alexa 488 (Invitrogen # A-11001, 1:400) at 4 °C overnight or over two nights (Fig. 2E-L). Brains were washed 3 times, 20 min each with 0.3% PBS-T followed by 2 quick washes with 1X PBS. The brains were mounted in Vectashield mounting medium (Vector Laboratories # H-1000-10) in SecureSeal imaging spacers (Electron Microscopy Sciences #70327-9S). Brains were imaged using Leica SP8 confocal microscope (Leica Microsystem) and deconvoluted using either Lightning Deconvolution (Leica Microsystem) or Huygens Confocal Deconvolution (Scientific Volume Imaging). The images were processed using Fiji ^72^.

### Total RNA isolation

Pupal brains were dissected in PBS and immediately frozen on dry ice. Totally, 30 brains per genotype were collected at 24h APF (After Puparium Formation). Three independent biological replicates per genotype were used for RNA isolation. Total RNA was isolated using RNeasy Plus Universal Kit (QIAGEN, USA) following the manufacturer’s instructions including DNase I treatment using RNase-Free DNase Set and on-column protocol (QIAGEN, USA). The isolated total RNAs were aliquoted for both RNA-seq and qRT-PCR.

### Quantitative reverse transcription PCR (RT-qPCR)

One microgram of total RNA from wild-type, Tet, and Fmr1 mutants was reserve transcribed to cDNA using MMLV High-Performance Reverse Transcriptase Kit (Biosearch Technologies #RT80125K) and oligo(dT) primers (IDT). The cDNAs were used for real-time quantitative PCR using gene-specific primers with Rpl32 as the internal reference control (**Supplementary Table S2**) and PowerUp SYBR Green Master Mix (Applied Biosystems #A25741) with StepOnePlus Real-Time PCR System (Applied Biosystems). The relative quantification (RQ) for the expression of each gene was calculated using the comparative Ct method ^73^.

### RNA-seq library generation

Library preparation and sequencing were performed at Novogene. One microgram of total RNA was used as input for library preparation. Prior to library preparation, total RNA quality was validated for the following parameters: degradation and contamination using 1% agarose gel electrophoresis, purity by NanoPhotometer® spectrophotometer (IMPLEN, USA), and integrity using Bioanalyzer 2100 and RNA Nano 6000 Kit (Agilent Technologies, USA). Sequencing libraries were generated using NEBNext® Ultra™ Directional RNA Library Prep Kit for Illumina® (NEB, USA) following the manufacturer’s instruction. mRNA was purified from total RNA using oligo dT magnetic beads. Purified mRNAs were fragmented by divalent cations. First-strand cDNA synthesis using random hexamer and M-MuLV RT (RNase H-). The dUTP method was used to synthesize the second strand and generate a strand-specific library. After 3’ A overhangs were cleaved by exonuclease, DNA fragments were ligated with NEBNext® hairpin loop adaptor to prepare for hybridization. AMPure XP system (Beckman Coulter, USA) was used for 150-200 bp size selection. The size selected and purified products were treated with USER® Enzyme to open up the NEBNext adapter. Then, PCR was performed with Phusion High-Fidelity DNA polymerase using Universal PCR primers and Index (X) prime. PCR products were purified using the AMPure XP system and quality was assessed by Agilent Bioanalyzer 2100. Clustering was done using cBot Cluster System using PE Cluster Kit cBot-HS (Illumina, USA) according to the manufacturer’s recommendation. Sequencing was performed on Illumina Novaseq 6000 (Illumina, USA) to generate paired-end reads.

### RNA-seq data analysis

Raw reads were filtered and trimmed by Trimmomatic 0.39 ^74^. Then, the filtered reads will be mapped to Ensembl *Drosophila melanogaster* genome assembly BDGP6.22 using HISAT2 ^75^. The mapping SAM files were converted to BAM using Samtools ^76^. The number of reads mapped to the individual gene is counted by featureCounts ^77^. Differential gene expression analysis was done by DESeq2 ^78^. Then, the result was visualized by the Bioconductor package EnhancedVolcano ^79^.

### Western blot

Thirty L3 instar larval brains from Act5C-GAL4 driving Gs2 CRISPR/Cas9 mutagenesis or Act5C- GAL4 wild-type control were dissected and immediately frozen on dry ice. Total protein was isolated from these brains using RIPA buffer (25 mM Tris HCl pH 7.6, 150 mM NaCl, 1% NP-40, 1% sodium deoxycholate, 0.1% SDS; Thermo Scientific #89900) supplemented with 1X Protease Inhibitor Cocktail (Roche # 5892791001) and 25ug of the total protein was loaded to each well. Glutamine synthetase antibody, clone GS-6 (Millipore-Sigma #MAB302) was used at 1: 2000 dilution and beta-tubulin antibody, clone TU27 (BioLegend # 903401) was used at 1: 1000 dilution. The western blot signals were detected using anti-mouse IRDye 800CW Infrared Dyes conjugated secondary antibody in the LICOR Odyssey CLx imaging system. Signals were quantified using LICOR Image Studio Lite software.

### MPEP and glutamate treatment

Wild type and Tet^AXXC^ mutant were grown in the standard medium at 25°C to early third instar larva. Then, sixty larvae were collected and transferred to a vial containing MPEP- or glutamate-supplemented medium at 100 µM or water-supplemented medium as diluent control. Twelve hours after transferring, larvae that came out from the medium will be eliminated to ensure all larvae will consume the supplemented medium for at least 12 hours. This allows the larvae to uptake MPEP or glutamate before they begin to pupate and stop eating. The larvae that stayed inside the medium were continuing to grow into adults. Then, the adult flies after eclosion were transferred back to the standard medium. Five to ten days old adult brains were dissected and immunostained to quantify the percentage of MB β lobe fusion per each condition.

### Generating UAS-Gs2 transgene stocks

Gs2 cDNA was obtained from DGRC clone LP04559 (DGRC #1337593). The coding region was amplified by Phusion Hot Start II HF (Thermo Scientific # F565S) using gene-specific primers with homology arms designed for DNA Assembly: Gs2.RB.armR.Eco (gccgcagatctgttaacgctattcgtccaggcagatggt) and Gs2.RB.armL.Eco (gaactctgaatagggaattgggcaacatgcattccgcgatcctggag). The amplicon was cloned to the pUASTattB expression vector (DGRC #1419, deposited by Bischof and Basler) at the EcoRI restriction site by NEBuilder HiFi DNA Assembly Cloning Kit (NEB #E5520S). The pUASTattB-Gs2 vector was injected to *y*^1^ *w*^67c23^*; PCaryPattP40* stock ^80^ by the BestGene company to insert the UAS-Gs2 construct to attP40 landing site on the second chromosome. The *UAS-Gs2* stock was screened by the mini-white marker and validated by genomic PCR and Sanger sequencing.

*UAS-GS2* stock was crossed with Tet^AXXC^ and Fmr1^3^ mutants to generate *UAS-Gs2; Tet^AXXC^/TM6B Tb^1^* and *UAS-Gs2; Fmr1*^3^*/TM6B Tb^1^*. The Tet^AXXC^ and Fmr1^3^ mutants were also crossed with the Dilp2- GAL4 driver to generate *Dilp2-GAL4; Tet^AXXC^/TM6 Sb^1^* and *Dilp2-GAL4; Fmr1^3^/TM6 Sb^1^*. Then, the stocks will be crossed to generate rescued stocks and respective controls. The progenies were collected after eclosion, separated to eliminate balancers, transferred to fresh medium, and dissected at five to ten days old for MB β lobe immunostaining and quantification.

### Phenotypic quantification and statistical analysis

MB axon guidance phenotype was quantified based on previous described method ^20, 41, 48^ with the following modification, the MB β lobe fusion phenotype was categorized into fusion and no fusion groups. The statistical analyses between Gs2 rescue and Tet or Fmr1 mutants were done using Chi-square tests, GraphPad Prism version 9.5.1 (GraphPad Software, San Diego, California USA).

### Data and codes availability

RNA-seq data are available at the NCBI GEO database with accession number GSE231534 and token ctifkiyivdcrnit. Codes used for RNA-seq data analysis is available on GitHub at https://github.com/hhiept/tetxglu.

## Competing interest statement

The authors have declared no competing interests.

## Author contributions

Conceptualization (H.T., R.S.), Data curation (H.T., L.L., B.N.S., J.K., R.S.), Formal analysis (H.T., R.S.), Funding acquisition (R.S.), Investigation (H.T., L.L., B.N.S., J.K.), Methodology (H.T., L.L., B.N.S., J.K., R.S.), Project administration (H.T., R.S.), Resources (H.T., L.L., B.N.S., J.K., R.S.), Software (H.T.), Supervision (R.S.), Validation (H.T., L.L., J.K., R.S.), Visualization (H.T.), Writing-original draft (H.T.), Writing-review & editing (H.T., J.K., R.S.).

## Acknowledgments

We thank Kenneth Irvine, Cordelia Rauskolb, and Nicholas Stavropoulos for helpful comments on the manuscript. We want to thank Fei Wang, Kishan Bulsara, Anna Zhang, Ethan Chiang for preliminary data. Irvine, Barber, and Stavropoulos labs for helpful discussion and technical support, and Thomas A. Jongens and Eric Rulifson for fly stocks. We thank Svetlana Minakhina for the co-IP result in SI Appendix Fig. S10. We also thank Drosophila Genomics Resource Center, Bloomington Drosophila Stock Center, Vienna Drosophila Resource Center, and Developmental Studies Hybridoma Bank for cDNA, fly stocks, and antibodies. This work was supported by grants from the National Institute of Health (R01 GM118404 to R.S.), the Vietnam Education Foundation (VEF) and Charles and Johanna Busch Pre-doctoral Fellowships (to H.T.).

## Notes

### Summary of Updates

Manuscript, Figures, and Supplementary data updated.

## References

1. Tahiliani, M., Koh, K.P., Shen, Y., Pastor, W.A., Bandukwala, H., Brudno, Y., Agarwal, S., Iyer, L.M., Liu, D.R., Aravind, L., and Rao, A. (2009). Conversion of 5-methylcytosine to 5- hydroxymethylcytosine in mammalian DNA by MLL partner TET1. Science 324, 930–935. 10.1126/science.1170116.

2. Fu, L., Guerrero, C.R., Zhong, N., Amato, N.J., Liu, Y., Liu, S., Cai, Q., Ji, D., Jin, S.G., Niedernhofer, L.J., et al. (2014). Tet-mediated formation of 5-hydroxymethylcytosine in RNA. J Am Chem Soc 136, 11582–11585. 10.1021/ja505305z.

3. Delatte, B., Wang, F., Ngoc, L.V., Collignon, E., Bonvin, E., Deplus, R., Calonne, E., Hassabi, B., Putmans, P., Awe, S., et al. (2016). RNA biochemistry. Transcriptome-wide distribution and function of RNA hydroxymethylcytosine. Science 351, 282–285. 10.1126/science.aac5253.

4. Dai, H.Q., Wang, B.A., Yang, L., Chen, J.J., Zhu, G.C., Sun, M.L., Ge, H., Wang, R., Chapman, D.L., Tang, F., et al. (2016). TET-mediated DNA demethylation controls gastrulation by regulating Lefty-Nodal signalling. Nature 538, 528–532. 10.1038/nature20095.

5. Wang, F., Minakhina, S., Tran, H., Changela, N., Kramer, J., and Steward, R. (2018). Tet protein function during Drosophila development. PLoS One 13, e0190367. 10.1371/journal.pone.0190367.

6. Melamed, P., Yosefzon, Y., David, C., Tsukerman, A., and Pnueli, L. (2018). Tet Enzymes, Variants, and Differential Effects on Function. Front Cell Dev Biol 6, 22. 10.3389/fcell.2018.00022.

7. Beck, D.B., Petracovici, A., He, C., Moore, H.W., Louie, R.J., Ansar, M., Douzgou, S., Sithambaram, S., Cottrell, T., Santos-Cortez, R.L.P., et al. (2020). Delineation of a Human Mendelian Disorder of the DNA Demethylation Machinery: TET3 Deficiency. Am J Hum Genet 106, 234–245. 10.1016/j.ajhg.2019.12.007.

8. Seyama, R., Tsuchida, N., Okada, Y., Sakata, S., Hamada, K., Azuma, Y., Hamanaka, K., Fujita, A., Koshimizu, E., Miyatake, S., et al. (2022). Two families with TET3-related disorder showing neurodevelopmental delay with craniofacial dysmorphisms. J Hum Genet 67, 157–164. 10.1038/s10038-021-00986-y.

9. Santos-Cortez, R.L.P., Khan, V., Khan, F.S., Mughal, Z.U., Chakchouk, I., Lee, K., Rasheed, M., Hamza, R., Acharya, A., Ullah, E., et al. (2018). Novel candidate genes and variants underlying autosomal recessive neurodevelopmental disorders with intellectual disability. Hum Genet 137, 735–752. 10.1007/s00439-018-1928-6.

10. Harripaul, R., Vasli, N., Mikhailov, A., Rafiq, M.A., Mittal, K., Windpassinger, C., Sheikh, T.I., Noor, A., Mahmood, H., Downey, S., et al. (2018). Mapping autosomal recessive intellectual disability: combined microarray and exome sequencing identifies 26 novel candidate genes in 192 consanguineous families. Mol Psychiatry 23, 973–984. 10.1038/mp.2017.60.

11. Engle, E.C. (2010). Human genetic disorders of axon guidance. Cold Spring Harb Perspect Biol 2, a001784. 10.1101/cshperspect.a001784.

12. Strausfeld, N.J., Hansen, L., Li, Y., Gomez, R.S., and Ito, K. (1998). Evolution, discovery, and interpretations of arthropod mushroom bodies. Learn Mem 5, 11–37.

13. Aso, Y., Sitaraman, D., Ichinose, T., Kaun, K.R., Vogt, K., Belliart-Guerin, G., Placais, P.Y., Robie, A.A., Yamagata, N., Schnaitmann, C., et al. (2014). Mushroom body output neurons encode valence and guide memory-based action selection in Drosophila. Elife 3, e04580. 10.7554/eLife.04580.

14. Li, Q., Jang, H., Lim, K.Y., Lessing, A., and Stavropoulos, N. (2021). insomniac links the development and function of a sleep-regulatory circuit. Elife 10. 10.7554/eLife.65437.

15. Kunz, T., Kraft, K.F., Technau, G.M., and Urbach, R. (2012). Origin of Drosophila mushroom body neuroblasts and generation of divergent embryonic lineages. Development 139, 2510–2522. 10.1242/dev.077883.

16. Riedel, G., Platt, B., and Micheau, J. (2003). Glutamate receptor function in learning and memory. Behav Brain Res 140, 1–47. 10.1016/s0166-4328(02)00272-3.

17. Mattson, M.P. (2008). Glutamate and neurotrophic factors in neuronal plasticity and disease. Ann N Y Acad Sci 1144, 97–112. 10.1196/annals.1418.005.

18. Lujan, R., Shigemoto, R., and Lopez-Bendito, G. (2005). Glutamate and GABA receptor signalling in the developing brain. Neuroscience 130, 567–580. 10.1016/j.neuroscience.2004.09.042.

19. Wang, P.Y., Petralia, R.S., Wang, Y.X., Wenthold, R.J., and Brenowitz, S.D. (2011). Functional NMDA receptors at axonal growth cones of young hippocampal neurons. J Neurosci 31, 9289–9297. 10.1523/jneurosci.5639-10.2011.

20. McBride, S.M., Choi, C.H., Wang, Y., Liebelt, D., Braunstein, E., Ferreiro, D., Sehgal, A., Siwicki, K.K., Dockendorff, T.C., Nguyen, H.T., et al. (2005). Pharmacological rescue of synaptic plasticity, courtship behavior, and mushroom body defects in a Drosophila model of fragile X syndrome. Neuron 45, 753–764. 10.1016/j.neuron.2005.01.038.

21. Schmitz, Y., Luccarelli, J., Kim, M., Wang, M., and Sulzer, D. (2009). Glutamate controls growth rate and branching of dopaminergic axons. J Neurosci 29, 11973–11981. 10.1523/jneurosci.2927-09.2009.

22. Kreibich, T.A., Chalasani, S.H., and Raper, J.A. (2004). The neurotransmitter glutamate reduces axonal responsiveness to multiple repellents through the activation of metabotropic glutamate receptor 1. J Neurosci 24, 7085–7095. 10.1523/JNEUROSCI.0349-04.2004.

23. Santoro, M.R., Bray, S.M., and Warren, S.T. (2012). Molecular mechanisms of fragile X syndrome: a twenty-year perspective. Annu Rev Pathol 7, 219–245. 10.1146/annurev-pathol-011811-132457.

24. Dockendorff, T.C., Su, H.S., McBride, S.M., Yang, Z., Choi, C.H., Siwicki, K.K., Sehgal, A., and Jongens, T.A. (2002). Drosophila lacking dfmr1 activity show defects in circadian output and fail to maintain courtship interest. Neuron 34, 973–984. 10.1016/s0896-6273(02)00724-9.

25. Zhang, Y.Q., Bailey, A.M., Matthies, H.J., Renden, R.B., Smith, M.A., Speese, S.D., Rubin, G.M., and Broadie, K. (2001). Drosophila fragile X-related gene regulates the MAP1B homolog Futsch to control synaptic structure and function. Cell 107, 591–603. 10.1016/s0092-8674(01)00589-x.

26. Bolduc, F.V., Bell, K., Cox, H., Broadie, K.S., and Tully, T. (2008). Excess protein synthesis in Drosophila fragile X mutants impairs long-term memory. Nat Neurosci 11, 1143–1145. 10.1038/nn.2175.

27. Redt-Clouet, C., Trannoy, S., Boulanger, A., Tokmatcheva, E., Savvateeva-Popova, E., Parmentier, M.L., Preat, T., and Dura, J.M. (2012). Mushroom body neuronal remodelling is necessary for short-term but not for long-term courtship memory in Drosophila. Eur J Neurosci 35, 1684–1691. 10.1111/j.1460-9568.2012.08103.x.

28. Rasmussen, K.D., and Helin, K. (2016). Role of TET enzymes in DNA methylation, development, and cancer. Genes Dev 30, 733–750. 10.1101/gad.276568.115.

29. Williams, K., Christensen, J., Pedersen, M.T., Johansen, J.V., Cloos, P.A., Rappsilber, J., and Helin, K. (2011). TET1 and hydroxymethylcytosine in transcription and DNA methylation fidelity. Nature 473, 343–348. 10.1038/nature10066.

30. Deplus, R., Delatte, B., Schwinn, M.K., Defrance, M., Méndez, J., Murphy, N., Dawson, M.A., Volkmar, M., Putmans, P., Calonne, E., et al. (2013). TET2 and TET3 regulate GlcNAcylation and H3K4 methylation through OGT and SET1/COMPASS. Embo j 32, 645–655. 10.1038/emboj.2012.357.

31. Gratz, S.J., Wildonger, J., Harrison, M.M., and O’Connor-Giles, K.M. (2013). CRISPR/Cas9- mediated genome engineering and the promise of designer flies on demand. Fly (Austin) 7, 249–255. 10.4161/fly.26566.

32. Gratz, S.J., Ukken, F.P., Rubinstein, C.D., Thiede, G., Donohue, L.K., Cummings, A.M., and O’Connor-Giles, K.M. (2014). Highly specific and efficient CRISPR/Cas9-catalyzed homology- directed repair in Drosophila. Genetics 196, 961–971. 10.1534/genetics.113.160713.

33. Gratz, S.J., Cummings, A.M., Nguyen, J.N., Hamm, D.C., Donohue, L.K., Harrison, M.M., Wildonger, J., and O’Connor-Giles, K.M. (2013). Genome engineering of Drosophila with the CRISPR RNA-guided Cas9 nuclease. Genetics 194, 1029–1035. 10.1534/genetics.113.152710.

34. Lee, J.H., Voo, K.S., and Skalnik, D.G. (2001). Identification and characterization of the DNA binding domain of CpG-binding protein. J Biol Chem 276, 44669–44676. 10.1074/jbc.M107179200.

35. Nern, A., Pfeiffer, B.D., and Rubin, G.M. (2015). Optimized tools for multicolor stochastic labeling reveal diverse stereotyped cell arrangements in the fly visual system. Proc Natl Acad Sci U S A 112, E2967–2976. 10.1073/pnas.1506763112.

36. Diao, F., Ironfield, H., Luan, H., Diao, F., Shropshire, W.C., Ewer, J., Marr, E., Potter, C.J., Landgraf, M., and White, B.H. (2015). Plug-and-play genetic access to drosophila cell types using exchangeable exon cassettes. Cell Rep 10, 1410–1421. 10.1016/j.celrep.2015.01.059.

37. Lee, P.T., Zirin, J., Kanca, O., Lin, W.W., Schulze, K.L., Li-Kroeger, D., Tao, R., Devereaux, C., Hu, Y., Chung, V., et al. (2018). A gene-specific T2A-GAL4 library for Drosophila. Elife 7. 10.7554/eLife.35574.

38. Caggese, C., Barsanti, P., Viggiano, L., Bozzetti, M.P., and Caizzi, R. (1994). Genetic, molecular and developmental analysis of the glutamine synthetase isozymes of Drosophila melanogaster. Genetica 94, 275–281. 10.1007/bf01443441.

39. Thimgan, M.S., Berg, J.S., and Stuart, A.E. (2006). Comparative sequence analysis and tissue localization of members of the SLC6 family of transporters in adult Drosophila melanogaster. J Exp Biol 209, 3383–3404. 10.1242/jeb.02328.

40. Chaturvedi, R., Stork, T., Yuan, C., Freeman, M.R., and Emery, P. (2022). Astrocytic GABA transporter controls sleep by modulating GABAergic signaling in Drosophila circadian neurons. Curr Biol 32, 1895–1908.e1895. 10.1016/j.cub.2022.02.066.

41. Michel, C.I., Kraft, R., and Restifo, L.L. (2004). Defective neuronal development in the mushroom bodies of Drosophila fragile X mental retardation 1 mutants. J Neurosci 24, 5798–5809. 10.1523/jneurosci.1102-04.2004.

42. Eins, S., Spoerri, P.E., and Heyder, E. (1983). GABA or sodium-bromide-induced plasticity of neurites of mouse neuroblastoma cells in culture. A quantitative study. Cell Tissue Res 229, 457–460. 10.1007/bf00214987.

43. Barbin, G., Pollard, H., Gaïarsa, J.L., and Ben-Ari, Y. (1993). Involvement of GABAA receptors in the outgrowth of cultured hippocampal neurons. Neurosci Lett 152, 150–154. 10.1016/0304-3940(93)90505-f.

44. Sinakevitch, I., Grau, Y., Strausfeld, N.J., and Birman, S. (2010). Dynamics of glutamatergic signaling in the mushroom body of young adult Drosophila. Neural Dev 5, 10. 10.1186/1749-8104-5-10.

45. Enell, L.E., Kapan, N., Söderberg, J.A., Kahsai, L., and Nässel, D.R. (2010). Insulin signaling, lifespan and stress resistance are modulated by metabotropic GABA receptors on insulin producing cells in the brain of Drosophila. PLoS One 5, e15780. 10.1371/journal.pone.0015780.

46. Cao, J., Ni, J., Ma, W., Shiu, V., Milla, L.A., Park, S., Spletter, M.L., Tang, S., Zhang, J., Wei, X., et al. (2014). Insight into insulin secretion from transcriptome and genetic analysis of insulin- producing cells of Drosophila. Genetics 197, 175–192. 10.1534/genetics.113.160663.

47. Bear, M.F., Huber, K.M., and Warren, S.T. (2004). The mGluR theory of fragile X mental retardation. Trends Neurosci 27, 370–377. 10.1016/j.tins.2004.04.009.

48. Chang, S., Bray, S.M., Li, Z., Zarnescu, D.C., He, C., Jin, P., and Warren, S.T. (2008). Identification of small molecules rescuing fragile X syndrome phenotypes in Drosophila. Nat Chem Biol 4, 256–263. 10.1038/nchembio.78.

49. MacArthur, I.C., and Dawlaty, M.M. (2021). TET Enzymes and 5-Hydroxymethylcytosine in Neural Progenitor Cell Biology and Neurodevelopment. Front Cell Dev Biol 9, 645335. 10.3389/fcell.2021.645335.

50. Weng, Y.L., An, R., Cassin, J., Joseph, J., Mi, R., Wang, C., Zhong, C., Jin, S.G., Pfeifer, G.P., Bellacosa, A., et al. (2017). An Intrinsic Epigenetic Barrier for Functional Axon Regeneration. Neuron 94, 337–346.e336. 10.1016/j.neuron.2017.03.034.

51. Ismail, J.N., Badini, S., Frey, F., Abou-Kheir, W., and Shirinian, M. (2019). Drosophila Tet Is Expressed in Midline Glia and Is Required for Proper Axonal Development. Front Cell Neurosci 13, 252. 10.3389/fncel.2019.00252.

52. Frey, F., Sandakly, J., Ghannam, M., Doueiry, C., Hugosson, F., Berlandi, J., Ismail, J.N., Gayden, T., Hasselblatt, M., Jabado, N., and Shirinian, M. (2022). Drosophila Tet Is Required for Maintaining Glial Homeostasis in Developing and Adult Fly Brains. eNeuro 9. 10.1523/eneuro.0418-21.2022.

53. Singh, B.N., Tran, H., Kramer, J., Kirichenko, E., Changela, N., Wang, F., Feng, Y., Kumar, D., Lan, J., Bizet, M., et al. (2023). Tet-dependent 5-hydroxymethyl-Cytosine modification of mRNA regulates axon guidance genes in Drosophila. bioRxiv. 10.1101/2023.01.03.522592.

54. Temme, C., Zhang, L., Kremmer, E., Ihling, C., Chartier, A., Sinz, A., Simonelig, M., and Wahle, E. (2010). Subunits of the Drosophila CCR4-NOT complex and their roles in mRNA deadenylation. RNA 16, 1356–1370. 10.1261/rna.2145110.

55. Collart, M.A. (2016). The Ccr4-Not complex is a key regulator of eukaryotic gene expression. Wiley Interdiscip Rev RNA 7, 438–454. 10.1002/wrna.1332.

56. Tan, W., Schauder, C., Naryshkina, T., Minakhina, S., and Steward, R. (2016). Zfrp8 forms a complex with fragile-X mental retardation protein and regulates its localization and function. Dev Biol 410, 202–212. 10.1016/j.ydbio.2015.12.008.

57. Rulifson, E.J., Kim, S.K., and Nusse, R. (2002). Ablation of insulin-producing neurons in flies: growth and diabetic phenotypes. Science 296, 1118–1120. 10.1126/science.1070058.

58. Nässel, D.R. (2012). Insulin-producing cells and their regulation in physiology and behavior of Drosophila. Canadian Journal of Zoology 90, 476–488. 10.1139/z2012-009.

59. Barber, A.F., Erion, R., Holmes, T.C., and Sehgal, A. (2016). Circadian and feeding cues integrate to drive rhythms of physiology in Drosophila insulin-producing cells. Genes Dev 30, 2596–2606. 10.1101/gad.288258.116.

60. Nässel, D.R., Kubrak, O.I., Liu, Y., Luo, J., and Lushchak, O.V. (2013). Factors that regulate insulin producing cells and their output in Drosophila. Front Physiol 4, 252. 10.3389/fphys.2013.00252.

61. Singh, B.N., Tran, H., Kramer, J., Kirishenko, E., Changela, N., Wang, F., Feng, Y., Kumar, D., Tu, M., Liang, S., et al. (2023). Tet-dependent 5-hydroxymethyl-Cytosine modification of mRNA regulates the axon guidance genes robo2 and slit in Drosophila. bioRxiv. 10.1101/2023.01.03.522592.

62. Featherstone, D.E., Rushton, E., and Broadie, K. (2002). Developmental regulation of glutamate receptor field size by nonvesicular glutamate release. Nat Neurosci 5, 141–146. 10.1038/nn789.

63. Martinez-Hernandez, A., Bell, K.P., and Norenberg, M.D. (1977). Glutamine synthetase: glial localization in brain. Science 195, 1356–1358. 10.1126/science.14400.

64. Feldmann, N., del Rio, R.M., Gjinovci, A., Tamarit-Rodriguez, J., Wollheim, C.B., and Wiederkehr, A. (2011). Reduction of plasma membrane glutamate transport potentiates insulin but not glucagon secretion in pancreatic islet cells. Mol Cell Endocrinol 338, 46–57. 10.1016/j.mce.2011.02.019.

65. Anderson, C.M., Bridges, R.J., Chamberlin, A.R., Shimamoto, K., Yasuda-Kamatani, Y., and Swanson, R.A. (2001). Differing effects of substrate and non-substrate transport inhibitors on glutamate uptake reversal. J Neurochem 79, 1207–1216. 10.1046/j.1471-4159.2001.00668.x.

66. Prickett, T.D., and Samuels, Y. (2012). Molecular pathways: dysregulated glutamatergic signaling pathways in cancer. Clin Cancer Res 18, 4240–4246. 10.1158/1078-0432.Ccr-11-1217.

67. Miladinovic, T., Nashed, M.G., and Singh, G. (2015). Overview of Glutamatergic Dysregulation in Central Pathologies. Biomolecules 5, 3112–3141. 10.3390/biom5043112.

68. Montanari, M., Martella, G., Bonsi, P., and Meringolo, M. (2022). Autism Spectrum Disorder: Focus on Glutamatergic Neurotransmission. Int J Mol Sci 23. 10.3390/ijms23073861.

69. Strauss, A.L., Kawasaki, F., and Ordway, R.W. (2015). A Distinct Perisynaptic Glial Cell Type Forms Tripartite Neuromuscular Synapses in the Drosophila Adult. PLoS One 10, e0129957. 10.1371/journal.pone.0129957.

70. Viswanathan, S., Williams, M.E., Bloss, E.B., Stasevich, T.J., Speer, C.M., Nern, A., Pfeiffer, B.D., Hooks, B.M., Li, W.P., English, B.P., et al. (2015). High-performance probes for light and electron microscopy. Nat Methods 12, 568–576. 10.1038/nmeth.3365.

71. Tran, H.H., Dang, S.N.A., Nguyen, T.T., Huynh, A.M., Dao, L.M., Kamei, K., Yamaguchi, M., and Dang, T.T.P. (2018). Drosophila Ubiquitin C-Terminal Hydrolase Knockdown Model of Parkinson’s Disease. Sci Rep 8, 4468. 10.1038/s41598-018-22804-w.

72. Schindelin, J., Arganda-Carreras, I., Frise, E., Kaynig, V., Longair, M., Pietzsch, T., Preibisch, S., Rueden, C., Saalfeld, S., Schmid, B., et al. (2012). Fiji: an open-source platform for biological-image analysis. Nat Methods 9, 676–682. 10.1038/nmeth.2019.

73. Livak, K.J., and Schmittgen, T.D. (2001). Analysis of relative gene expression data using real- time quantitative PCR and the 2(-Delta Delta C(T)) Method. Methods 25, 402–408. 10.1006/meth.2001.1262.

74. Bolger, A.M., Lohse, M., and Usadel, B. (2014). Trimmomatic: a flexible trimmer for Illumina sequence data. Bioinformatics 30, 2114–2120. 10.1093/bioinformatics/btu170.

75. Kim, D., Paggi, J.M., Park, C., Bennett, C., and Salzberg, S.L. (2019). Graph-based genome alignment and genotyping with HISAT2 and HISAT-genotype. Nat Biotechnol 37, 907–915. 10.1038/s41587-019-0201-4.

76. Li, H., Handsaker, B., Wysoker, A., Fennell, T., Ruan, J., Homer, N., Marth, G., Abecasis, G., and Durbin, R. (2009). The Sequence Alignment/Map format and SAMtools. Bioinformatics 25, 2078–2079. 10.1093/bioinformatics/btp352.

77. Liao, Y., Smyth, G.K., and Shi, W. (2019). The R package Rsubread is easier, faster, cheaper and better for alignment and quantification of RNA sequencing reads. Nucleic Acids Res 47, e47. 10.1093/nar/gkz114.

78. Love, M.I., Huber, W., and Anders, S. (2014). Moderated estimation of fold change and dispersion for RNA-seq data with DESeq2. Genome Biol 15, 550. 10.1186/s13059-014-0550-8.

79. Blighe, K., Rana, S., Turkes, E., Ostendorf, B., Grioni, A., and Lewis, M. (2022). EnhancedVolcano: Publication-ready volcano plots with enhanced colouring and labeling. R package version 1.16.0. R package version 1.16.0. 10.18129/B9.bioc.EnhancedVolcano

80. Markstein, M., Pitsouli, C., Villalta, C., Celniker, S.E., and Perrimon, N. (2008). Exploiting position effects and the gypsy retrovirus insulator to engineer precisely expressed transgenes. Nat Genet 40, 476–483. 10.1038/ng.101.

