## Supplementary information for "Tet Controls Axon Guidance in Early Brain Development through Glutamatergic Signaling"

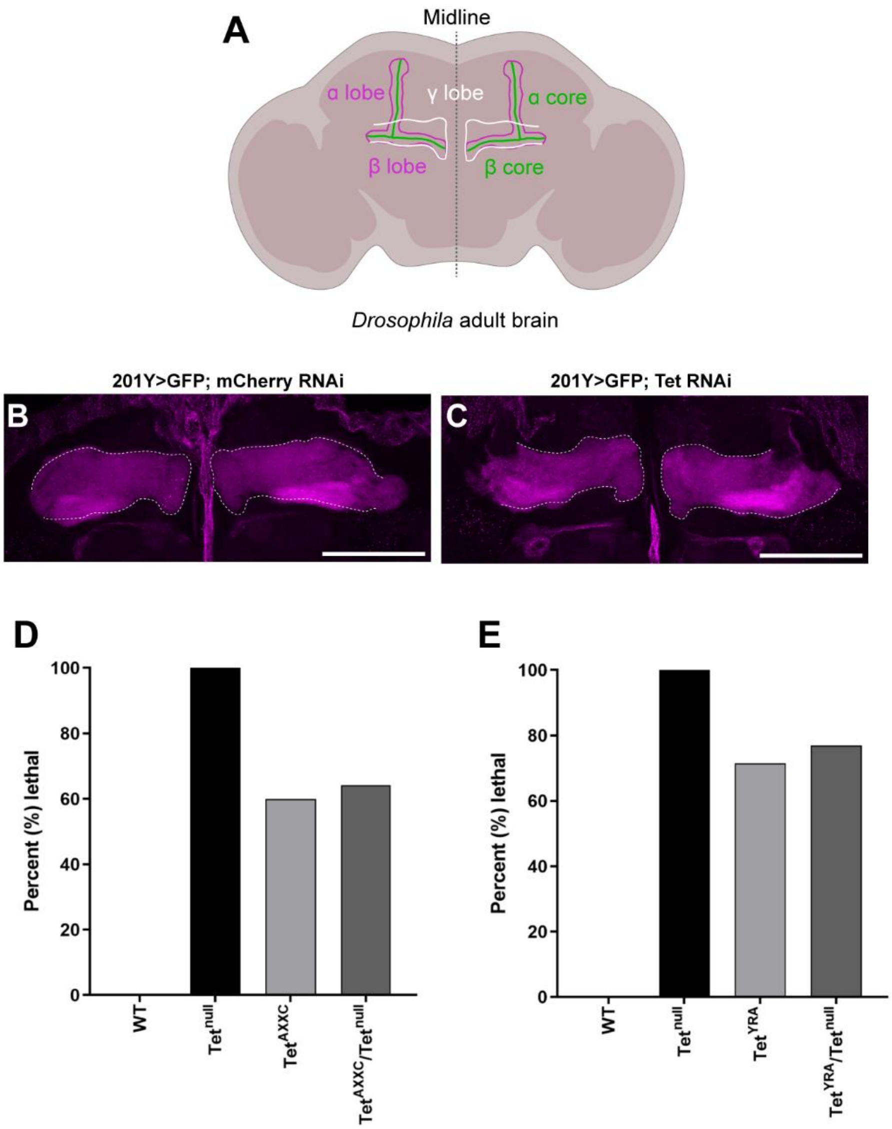


**Figure S1. Knockdown Tet using 201Y-GAL4 does not affect the MB γ axon guidance; Tet$^{AXXC}$ and Tet$^{YRA}$ exhibit partial lethality while Tet$^{null}$ is completely lethal at late pupa compared to the wild type.** (**A**) Illustration of 201Y>CD8::GFP expression pattern in the mushroom body lobes (green) and position of the mushroom body α, β (magenta), and γ (white) lobes in the adult brain. (**B**) 201Y> mCherry RNAi control and (**C**) 201Y>Tet RNAi knockdown. Adult brains were stained with anti-Fas2 antibody and sectioned by confocal to visualize γ lobes. Scale bar: 50 µm. (**D**) Homozygous Tet$^{AXXC}$ or Tet$^{AXXC}$/Tet$^{null}$ percent of survival compared to wild type and homozygous Tet$^{null}$. (**E**) Homozygous Tet$^{YRA}$ or Tet$^{YRA}$/Tet$^{null}$ percent of survival compared to wild type and homozygous Tet$^{null}$.

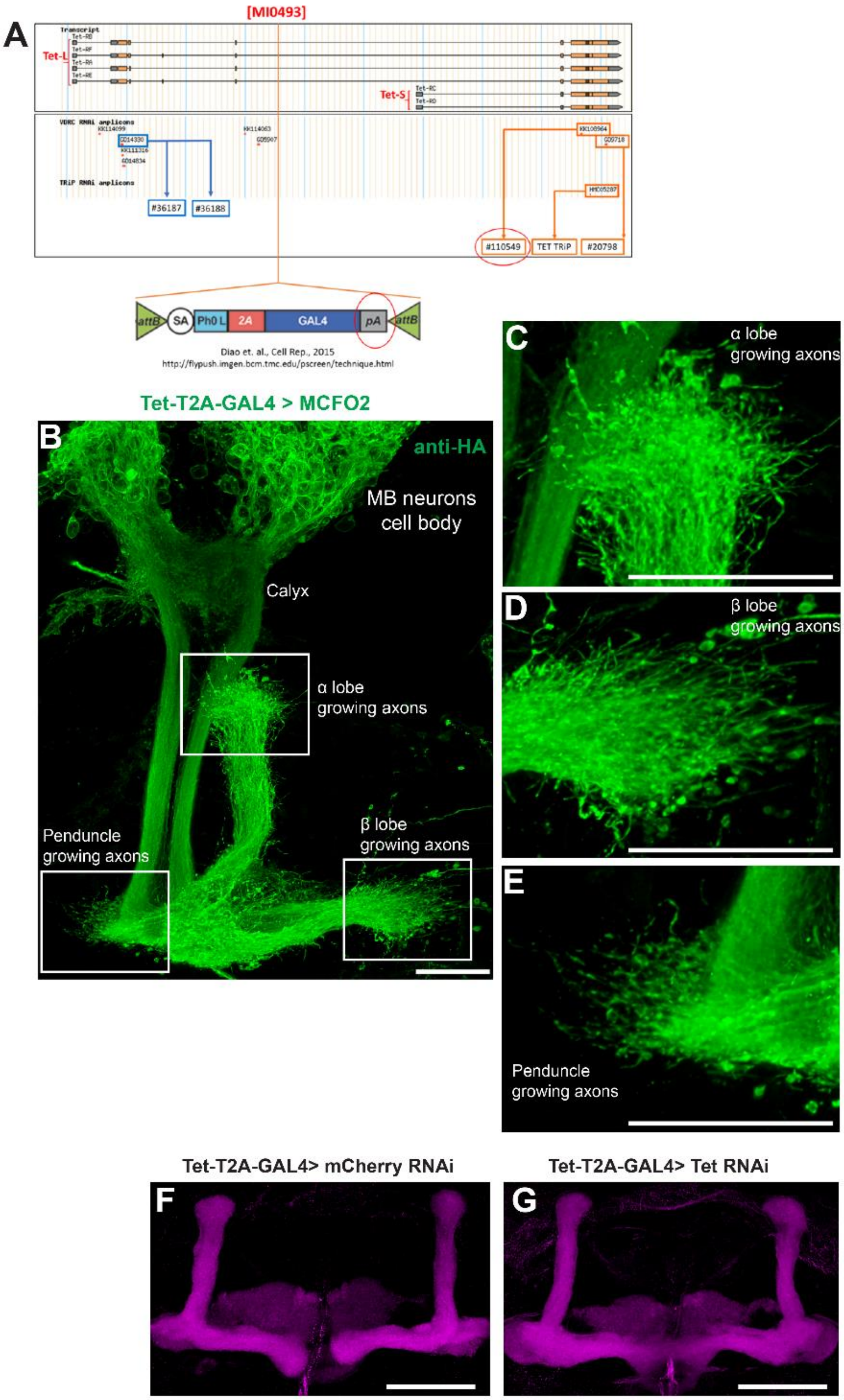


**Figure S2. Utilizing Tet-T2A-GAL4 to simultaneously knockdown Tet and labeling MB growing axons.** (**A**) Schematic representation of the site where Tet-T2A-GAL4 is inserted in the Tet gene [1] and the site where Tet RNAi #110549 (KK#108964) was designed to knockdown Tet. The poly(A) (pA) signal in the Tet-T2A-GAL4 construct will trigger transcriptional termination [2]; hence, the mRNA encoding for the GAL4 protein will not have the recognition site of Tet RNAi #110549 preventing it from being degraded by the RNAi mechanism when using Tet-T2A-GAL4 to drive Tet RNAi knockdown. (**B-E**) Tet-T2A-GAL4 expresses in MB neurons and can be used to label MB growing axons at early developing brain (**B**) The mushroom body neurons and their axons are labeled by Tet-T2A-GAL4 driven smGdP molecular markers at 21h APF brain (stained with HA antibody, green). (**C-E**) High-magnified insects from (B) show growing axons in α and β lobes, as well as the peduncle region with growth cones, projecting at the tips of each lobe or peduncle; scale bars: 25 µm. (**F-G**) Knockdown Tet using Tet-T2A-GAL4 driver produces same β lobe fusion phenotype as seen in Tet knockdown using 201Y-GAL4 driver. Tet-T2A-GAL4 drives mCherry RNAi control (**F**) or Tet RNAi knockdown (**G**); scale bar: 50 µm.

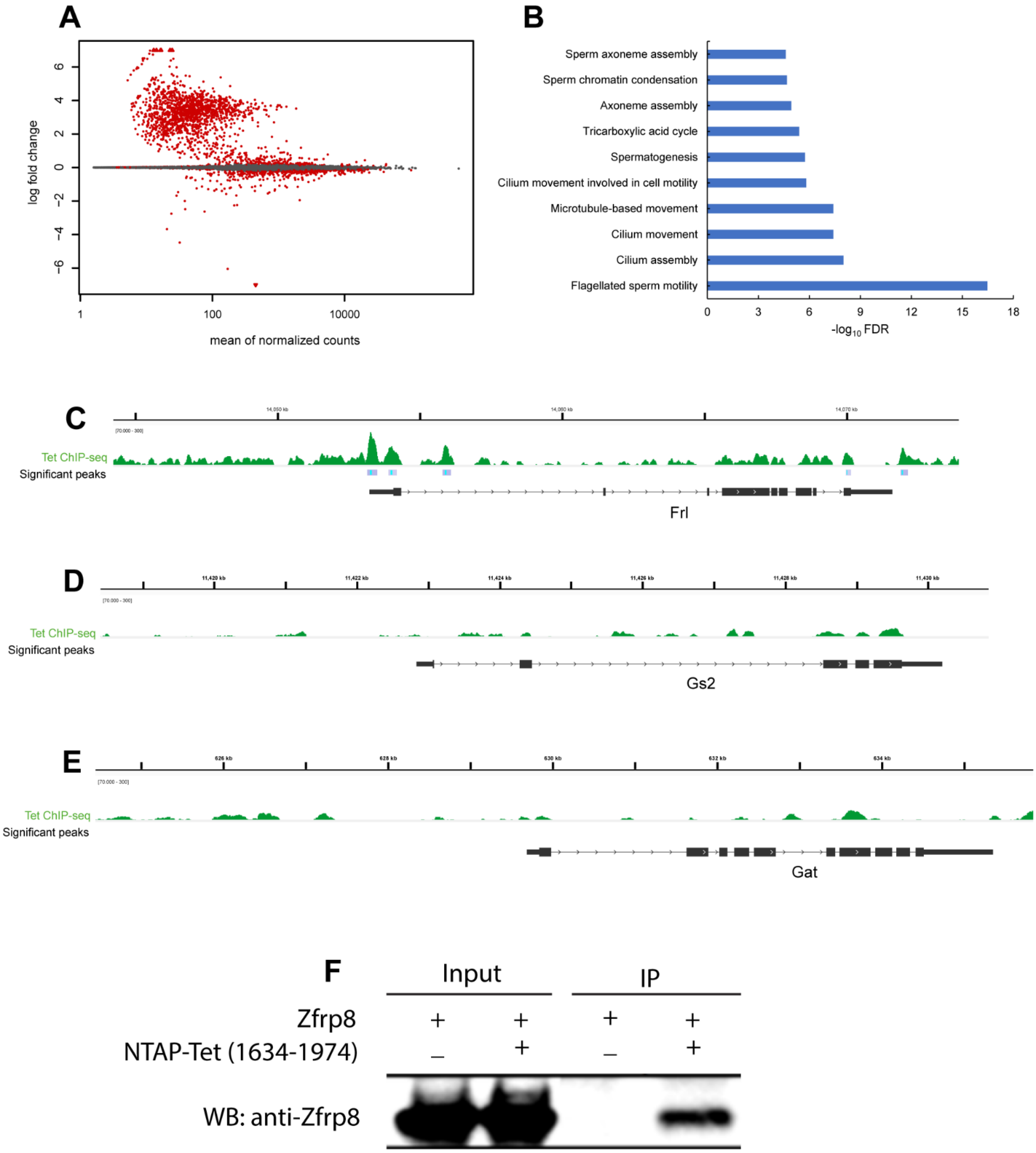


**Figure S3. Tet$^{AXXC}$ transcriptomic analysis and supporting data for how Tet regulates Gs2 and Gat. (A-B)** MA plot and GO term analysis of the differential expressed genes between Tet$^{AXXC}$ mutant and wild type. (**A**) MA plot showed a majority of up-regulated genes in Tet$^{AXXC}$ are low-expressed genes with around 100 read counts or less. (**B**) The top 10 GO terms of the up-regulated genes in Tet$^{AXXC}$ compared to wild type. (**C-E**) Tet binding visualized by Tet ChIP-seq data on three genes *Frl*, *Gs2*, and *Gat*. While Tet showed significant peaks in *Frl* gene (**C**) according to your recent study [3], there are no significant binding peaks of Tet on *Gs2* (**D**) and *Gat* (**E**) genes, and the extended regions from the 5' end of the genes. (**F**) Zfrp8 co-immunoprecipitated with NTAP tagged *Drosophila* Tet (aa 1634-1974) expressed in S2 cells.

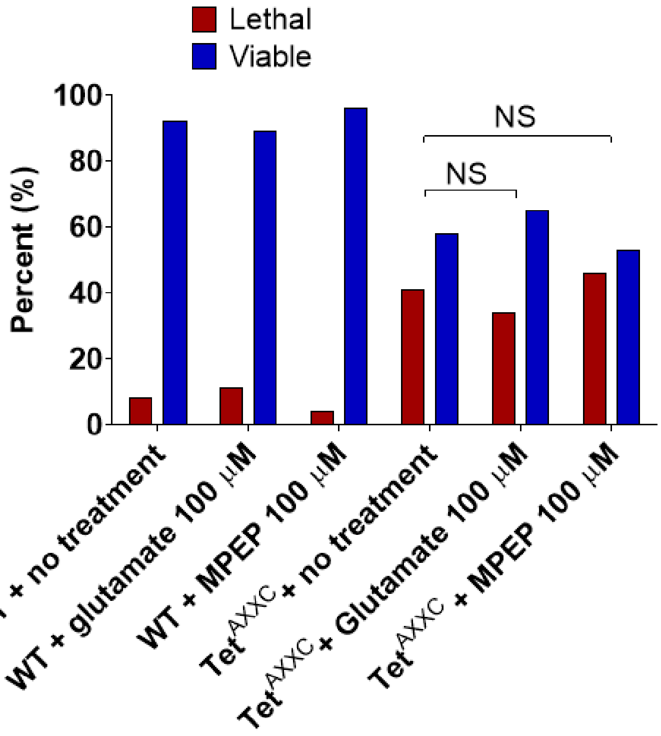


**Figure S4. MPEP and glutamate treatment do not affect the viability of both wild type and Tet$^{AXXC}$ mutant.** Percentage of viable (blue) or lethal (red) pupae obtained after the treatments. There are no significant (NS) differences between MPEP or glutamate-treated group versus the no-treatment group; Chi-square test with $p > 0.05$.

**Supplementary Table S1.** sgRNA and primer sequences for CRISPR/Cas9 and HDR mediated *Tet*$^{AXXC}$ and *Tet*$^{YRA}$ mutations.

| **Primers/gRNAs** | **Sequence** |
|---|---|
| gRNA_AXXC_1F | cttcGGGTAGGTGATCGAAGGGTG |
| gRNA_AXXC_1R | aaacCACCCTTCGATCACCTACCC |
| gRNA_AXXC_2F | cttcGGTCTTGTCGTTTCGGCAGG |
| gRNA_AXXC_2R | aaacCCTGCCGAAACGACAAGACC |
| HT299 | GGAGACGTATATGGTCTTCTTTTCCACACCAATGCCTGTGCCCATAG |
| HT300 | CAATTTTACGCAGACTATCTTTCTAGGGTTAATCGCAGCGGTTGCCAGTTC |
| HT293 | CACAATATGATTATCTTTCTAGGGTTAACAGGACTACGGTCGTCGCCACC |
| HT290 | GGCCCGCTTGCGCTTCTTCTTGGAC |
| HT298 | CCAAGAAGAAGCGCAAGCGGGCCGGCGAATGCGTGGGATGC |
| HT303 | GGAGACCTATAGTGTCTTCGGGGAGATCCCAGATTCGGGAACCTTTCAC |
| gRNA_YRA_1F | CTTCGCTGTCCGTAAAGAGCGAGG |
| gRNA_YRA_1R | AAACCCTCGCTCTTTACGGACAGC |
| gRNA_YRA_2F | CTTCGCCTCGCTCTTTACGGACAGC |
| gRNA_YRA_2R | AAACGCTGTCCGTAAAGAGCGAGGC |

**Supplementary Table S2.** Primer sequences for RT-qPCR.

| **Primers** | **Sequence** |
|---|---|
| Gs2.FWD2 | GGATGGCCCGTTTCCTCT |
| Gs2.REV2 | ACCAGCACCGTTCCAATC |
| Gat.FWD1 | CGCAGAAATAGACCCAGGATGT |
| Gat.RWD1 | GCCGTTGACATGTTGAAACCAT |
| Rpl32.FWD | CATACAGGCCCAAGATCGTG |
| Rpl32.REV | GGCGACGCACTCTGTTGTC |

**Supplementary data 1.** Down-regulated genes in *Tet*$^{AXXC}$ compared to *wild type*.

*Supplementary data 1.xlsx*

**Supplementary data 2.** Up-regulated genes in *Tet*$^{AXXC}$ compared to *wild type*.

*Supplementary data 2.xlsx*

### References


1. Lee, P.T., Zirin, J., Kanca, O., Lin, W.W., Schulze, K.L., Li-Kroeger, D., Tao, R., Devereaux, C., Hu, Y., Chung, V., et al. (2018). A gene-specific T2A-GAL4 library for Drosophila. Elife *7*. 10.7554/eLife.35574.
2. Diao, F., Ironfield, H., Luan, H., Diao, F., Shropshire, W.C., Ewer, J., Marr, E., Potter, C.J., Landgraf, M., and White, B.H. (2015). Plug-and-play genetic access to drosophila cell types using exchangeable exon cassettes. Cell Rep *10*, 1410-1421. 10.1016/j.celrep.2015.01.059.
3. Singh, B.N., Tran, H., Kramer, J., Kirishenko, E., Changela, N., Wang, F., Feng, Y., Kumar, D., Tu, M., Liang, S., et al. (2023). Tet-dependent 5-hydroxymethyl-Cytosine modification of mRNA regulates the axon guidance genes robo2 and slit in Drosophila. bioRxiv. 10.1101/2023.01.03.522592.
